# Phylogenomics offers resolution of major tunicate relationships

**DOI:** 10.1101/236505

**Authors:** Kevin M. Kocot, Michael G. Tassia, Kenneth M. Halanych, Billie J. Swalla

## Abstract

Tunicata, a diverse clade of approximately 3,000 described species of marine, filter-feeding chordates, is of great interest to researchers because tunicates are the closest living relatives of vertebrates and they facilitate comparative studies of our own biology. The group also includes numerous invasive species that cause considerable economic damage and some species of tunicates are edible. Despite their diversity and importance, relationships among major lineages of Tunicata are not completely resolved. Here, we supplemented public data with transcriptomes from seven species spanning the diversity of Tunicata and conducted phylogenomic analyses on data sets of up to 798 genes. Sensitivity analyses were employed to examine the influences of reducing compositional heterogeneity and branch-length heterogeneity. All analyses maximally supported a monophyletic Tunicata within Olfactores (Vertebrata + Tunicata). Within Tunicata, all analyses recovered Appendicularia sister to the rest of Tunicata and confirmed (with maximal support) that Thaliacea is nested within Ascidiacea. Stolidobranchia is the sister taxon to all other tunicates except Appendicularia. In most analyses, phlebobranch tunicates were recovered paraphyletic with respect to Aplousobranchia. Support for this topology varied but was strong in some cases. However, when only the 50 best genes based on compositional heterogeneity were analysed, we recovered Phlebobranchia and Aplousobranchia reciprocally monophyletic with strong support, consistent with most traditional morphology-based hypotheses. Examination of internode certainty also cast doubt on results of phlebobranch paraphyly, which may be due to limited taxon sampling. Taken together, these results provide a higher-level phylogenetic framework for our closest living invertebrate relatives.

## 1. Introduction

Tunicata (Lamarck, 1816) is a diverse clade of approximately 3,000 described species of marine, filter-feeding chordates (Shenkar and Swalla, 2011). This morphologically plastic group, which may be benthic or pelagic and solitary or colonial, has intrigued researchers for a number of reasons. Its position as the sister lineage of vertebrates (e.g., Bourlat et al., 2006; Delsuc et al., 2005, 2006) makes it an import resource for studying the evolution of developmental mechanisms (e.g., Racioppi et al., 2017; Suzuki et al., 2016; Swalla and Jeffery, 1996; Taketa and De Tomaso, 2015). Additionally, Tunicata includes some of the most invasive benthic marine animals known (e.g., Kaplan et al., 2017; Reem et al., 2013), which can cause catastrophic damage if left unchecked. Lastly, tunicates are eaten in some parts of the world, thus supporting fishery industries (Lambert et al., 2016). Despite their importance, evolutionary relationships within Tunicata remain uncertain, hindering understanding of the evolutionary history and ancestral character states of its constituent lineages and Chordata as a whole.

Tunicates date back to the Early Cambrian (Chen et al., 2003; Shu et al., 2001). Lamarck (1816) first described modern tunicates for their hard, leathery outer covering, the “tunic,” and included ascidians, pyrosomes and salps within the group. Tunicates were subsequently divided into three recognized classes: Ascidiacea (sea squirts), Thaliacea (pelagic salps, doliolids, and pyrosomes; Berrill, 1936), and Appendicularia (larvaceans), which have been associated with a suite of gross morphological and life history features (Table 1). Of these classes, Ascidiacea, comprised of sessile tunicates, is the most species rich (∼3,000 species), but a growing body of evidence indicates that the pelagic Thaliacea (∼100 species) is nested within Ascidiacea (Swalla et al., 2000; Stach et al., 2002; Tsagkogeorga et al., 2009; Winchell et al., 2002; Zeng et al., 2006).

**Table 1.** Gross morphology and life history features of major tunicate clades

| Taxon | Benthic/Pelagic | Solitary/Colonial | Branchial sac | Gonad position |
| --- | --- | --- | --- | --- |
| Appendicularia | Pelagic | Solitary | None | Dorsal |
| Phlebobranchia | Benthic | Usually solitary | Vascular | Enterogona<sup>1</sup> |
| Aplousobranchia | Benthic | Colonial | Simple | Enterogona<sup>1</sup> |
| Stolidobranchia | Benthic | Solitary or Colonial | Folded | Pleurogona<sup>2</sup> |
| Thaliacea | Pelagic | Solitary or Colonial | Simple | Enterogona<sup>1</sup> |

<sup>1</sup>Unpaired gonads are situated on one side or behind the intestinal loop. <sup>2</sup>Gonads are in the lateral mantle wall on both sides.

Appendicularia is a group of small, pelagic tunicates characterized by a putatively paedomorphic adult body plan that resembles the tadpole larva, very short generation times, and secretion of a unique mucous-like “house” used in feeding (Stach et al., 2008). Most appendicularian genes studied to date exhibit high rates of nucleotide substitution leading to long branches in molecular phylogenetic analyses (e.g., Delsuc et al., 2006, 2008; Govindarajan et al., 2011; Stach and Turbeville, 2002; Swalla et al., 2000; Wada, 1998). Previous studies have recovered appendicularians as either sister to the rest of Tunicata or within Ascidiacea (reviewed by Tsagkogeorga et al., 2009).

Higher-level taxonomy of Ascidiacea is based primarily on morphological characters of the branchial sac, the organ used to collect particles from the water column, and as such they are divided into three traditionally recognized orders: Aplousobranchia (simple branchial sac), Phlebobranchia (vascular branchial sac), and Stolidobranchia (folded branchial sac; Lahille, 1886). However, relationships among these clades remain uncertain and, as noted above, Thaliacea and Appendicularia may be nested within this clade of otherwise benthic tunicates. Stolidobranchia is a large and morphologically heterogeneous clade of solitary, social, and colonial tunicates (Zeng and Swalla, 2005; Zeng et al., 2006). Despite this, molecular phylogenetic analyses based on 18S rDNA and mitochondrial genes have shown convincingly that Molgulidae, a group of solitary ascidians with various larval morphologies, is sister to all other stolidobranchs whereas relationships within Pyuridae and Styelidae, the two other major lineages of Stolidobranchia, have generally not been well-resolved (e.g., Stach et al., 2002; Swalla et al., 2000; Tsagkogeorga et al., 2009; Winchell et al., 2002; Zeng and Swalla, 2005; Zeng et al. 2006). Aplousobranchia has been recovered as unambiguously monophyletic in most molecular analyses to date (e.g., Stach and Turbeville, 2002; Tsagkogeorga et al., 2009; Turon and López-Legentil, 2004), but these colonial tunicates tend to have high rates of nucleotide substitution and support for their position relative to other tunicates has generally been weak. Phlebobranchia is a traditionally recognized group of mostly solitary tunicates, but the composition of this clade has been debated. For example, Cionidae, which includes the widely-studied species *Ciona intestinalis, C. robusta* and *C. savignyi,* was originally included within Phlebobranchia (Berrill, 1936). However, this genus of important model tunicate species has also been viewed as a subclade of Aplousobranchia (e.g., Kott, 1990, 1969).

Although molecular phylogenetic studies of tunicates conducted to date (Govindarajan et al., 2011; Shenkar et al., 2016; Singh et al., 2009; Stach et al., 2010; Stach and Turbeville, 2002; Swalla et al., 2000; Tsagkogeorga et al., 2009; Turon and López-Legentil, 2004; Zeng et al., 2006; Zeng and Swalla, 2005) have greatly advanced understanding of relationships within some clades, tunicate higher-level phylogeny has been difficult to reconstruct. Evolutionary history of Tunicata has likely been a particularly challenging question in invertebrate systematics because several tunicate lineages (Appendicularia, Thaliacea, and many species within Aplousobranchia) exhibit long branch lengths for 18S and at least some other genes (Stach and Turbeville, 2002; Swalla et al., 2000; Tsagkogeorga et al., 2009; Winchell et al., 2002; Yokobori et al., 2006; Zeng et al., 2006).

Phylogenomic analyses have been important to our understanding of chordate evolutionary history by showing that tunicates and not cephalochordates are the sister taxon of the vertebrates (Bourlat et al., 2006; Delsuc et al. 2006, 2008; Dunn et al. 2008; Putnam et al. 2008), but no study to date has had the necessary taxon sampling to address long-standing questions about evolutionary relationships among the major lineages of tunicates. To this end, we supplemented publicly available tunicate and outgroup genome and transcriptome data with transcriptomes from taxa spanning the diversity of Tunicata and re-evaluated the higher-level evolutionary history of this important group.

## 2. Methods

We sampled publicly available and newly generated transcriptome and/or genome data from all extant tunicate orders with the exception of Doliolida, which was previously shown to be nested within the otherwise well-sampled taxon Thaliacea (Govindarajan et al., 2011). Available tunicate data were augmented with new transcriptomes from specimens collected from Antarctica, the Northeastern Pacific, and the Northwestern Atlantic (Table 2). With the exception of the unidentified *Ascidia* sp. (?) from Antarctica, all of the newly sequenced species are well-known species from their respective collection localities that were easily identified based on habitus and structure of the branchial basket and gut. RNA was extracted and purified from RNAlater-preserved or frozen tissue samples using the Omega Bio-Tek Mollusc RNA kit or the Qiagen RNeasy kit. In either case, an on-column DNAse digestion was performed. RNA concentration was measured using a Qubit (Thermo Fisher) with the RNA High Sensitivity kit, RNA purity was assessed by measuring the 260/280 nm absorbance ratio using a Nanodrop Lite (Thermo Fisher), and RNA integrity was evaluated using a 1% SB agarose gel or a TapeStation (Agilent). At least 1 μg of total RNA for each specimen was sent to Macrogen (Cambridge, MA, USA) for Illumina TruSeq RNA v2 library preparation (polyA enrichment) and sequencing on the Illumina HiSeq 2500 system using the HiSeq SBS V4 chemistry with 100 bp paired-end sequencing. For *Styela gibbsii,* two separate libraries from material from the same individual were prepared and sequenced separately, but the raw reads were combined prior to transcriptome assembly.

**Table 2.** Specimen collection data.

| Species | Collection locality | Latitude | Longitude | Tissue extracted |
| --- | --- | --- | --- | --- |
| *Ascidia* sp. | Ross Sea, Antarctica | 78°03'47.7"S | 169°59'28.1"W | Gonad |
| *Corella willmeriana* | Roche Harbor, San Juan Island, WA | 48°36'34.6"N | 123°09'18.8"W | Gonad and branchial sac |
| *Distaplia occidentalis* | Roche Harbor, San Juan Island, WA | 48°36'34.6"N | 123°09'18.8"W | Entire small colony |
| *Pyrosomella verticillata* | Northwestern Atlantic | 37°45'46.8"N | 73°38'37.8"W | Several entire zooids |
| *Salpa fusiformis* | Northwestern Atlantic | 39°15'07"N | 71°56'24"W | Gonad and branchial sac |
| *Styela gibbsii* | Friday Harbor, San Juan Island, WA | 48°32'43.2"N | 123°00'43.2"W | Gonad |
| *Iasis* (= *Weelia*) *cylindrica* | Northwestern Atlantic | 37°41'05"N | 73°37'07"W | Gonad and branchial sac |

Dataset processing followed the general approach of Kocot et al. (2017). Publicly available genomic data were downloaded as predicted proteins if available (Table 3). Otherwise, predicted transcripts from genomes or assembled transcriptomes were downloaded. Publicly available transcriptomes available only as raw read data and our new transcriptome data were assembled using Trinity 2.2.0 with the --trimmomatic and --normalize_reads flags (Grabherr et al., 2011). Transcripts were translated with TransDecoder 2.0.0 or 2.0.1 (Haas et al., 2014) using the UniProt SwissProt database (accessed on September 20^th^, 2016; The Uniprot Consortium, 2014) and PFAM (Pfam-A.hmm) version 27 (Finn et al., 2015).

**Table 3.**
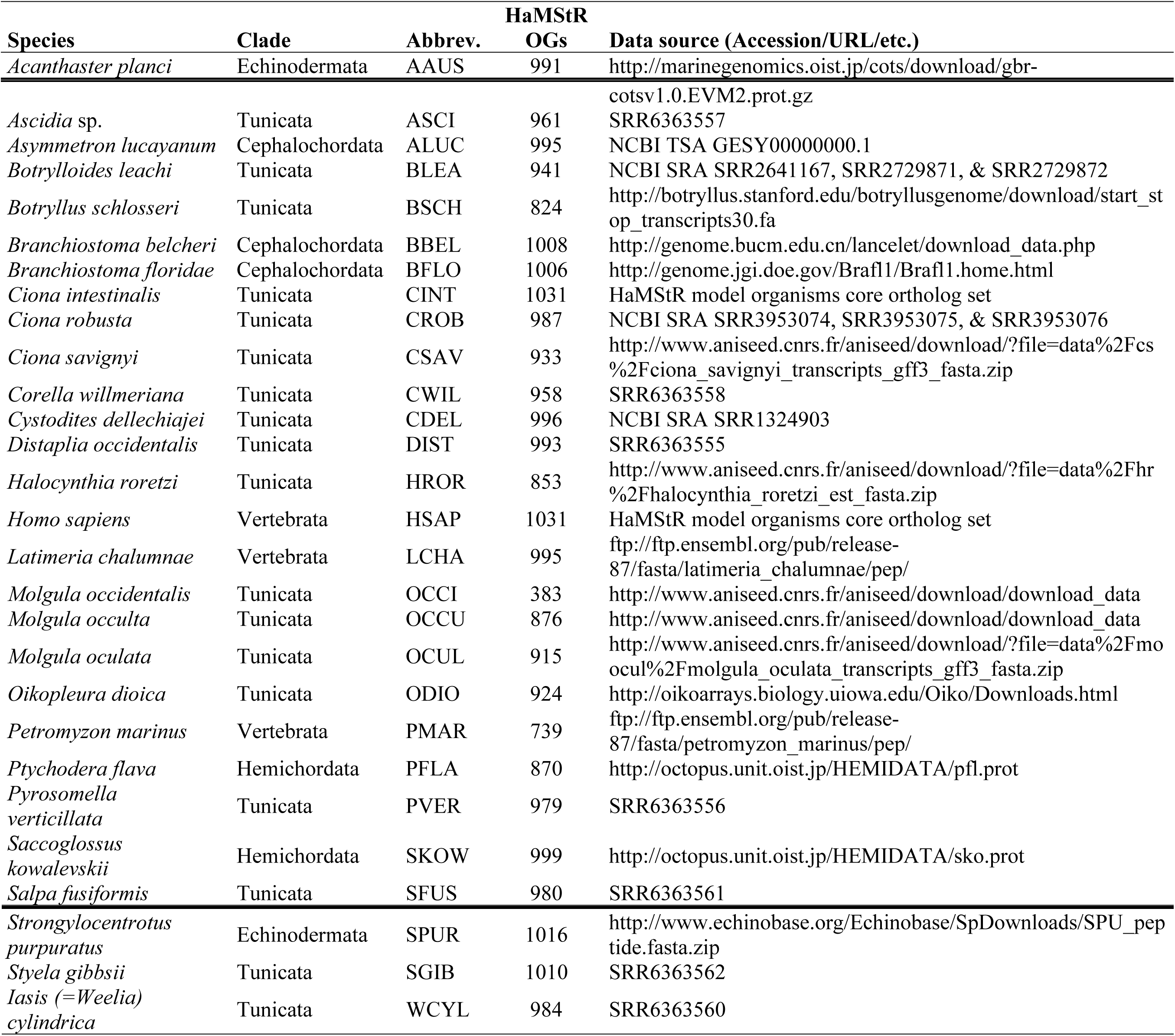
Taxon sampling, number of HaMStR orthologous groups (OGs) recovered for each taxon (out of 1,031), and sources of data used in phylogenomic analyses.

For orthology inference, we employed HaMStR 13 (Ebersberger et al., 2009) with the “model organsisms” core-ortholog set. Translated transcripts for all taxa except *Ciona intestinalis* and human were searched against the 1,031 profile hidden Markov models (pHMMs) using the default options. The “representative” flag was not used because it is not compatible with PhyloTreePruner (Kocot et al., 2013; see below). Sequences matching a pHMM were compared to the proteome of *Ciona* using BLASTP with the default search settings of HaMStR. If the *Ciona* amino acid sequence contributing to the pHMM was the best BLASTP hit in each of these back-BLASTs, the sequence was then assigned to that putative orthology group (simply referred to as “gene” henceforth). Redundant sequences that were identical (including partial sequences that were identical at least where they overlapped) were then removed with UniqHaplo (http://raven.iab.alaska.edu/∼ntakebay/), leaving only unique sequences for each taxon. Each gene was then aligned with MAFFT 7.273 using the automatic alignment strategy with a “maxiterate” value of 1,000 (Katoh and Standley, 2013). Alignments were then trimmed with Aliscore (Misof and Misof, 2009) and Alicut (Kück et al., 2010) with the default options to remove ambiguously aligned regions. Lastly, we deleted sequences that did not overlap with all other sequences in the alignment by at least 20 amino acids, starting with the shortest sequences not meeting this criterion. This step was necessary for downstream single-gene tree reconstruction. Finally, genes sampled for fewer than half of the 28 taxa after these steps were discarded.

In some cases, a taxon was represented in an alignment by two or more sequences (splice variants, lineage-specific gene duplications [=inparalogs], overlooked paralogs, or exogenous contamination). To screen for evidence of paralogy or contamination and select just one sequence for each taxon, an approximately maximum likelihood tree was inferred for each remaining alignment using FastTree 2 (Price et al., 2010) using the -slow and -gamma options. PhyloTreePruner (Kocot and Citarella et al., 2013) was then employed to use a tree-based approach to screen each single-gene alignment for evidence of paralogy or contamination. First, nodes with support values below 0.95 were collapsed into polytomies. Next, the maximally inclusive subtree was selected where each taxon was represented by no more than one sequence or, in cases where more than one sequence was present for any taxon, all sequences from that taxon formed a clade or were part of the same polytomy. Putative paralogs and contaminants (sequences falling outside of this maximally inclusive subtree) were then deleted from the input alignment. In cases where multiple sequences from the same taxon formed a clade or were part of the same polytomy, all sequences except the longest were deleted.

In order to further screen for genes or taxa with paralogy or contamination issues, genes that passed PhyloTreePruner screening and were still sampled for at least 15 of the 28 taxa were retained and used for single-gene tree building in RAxML 8.2.8 (Stamatakis, 2014) with the PROTGAMMAAUTOF model. The tree with the best likelihood score after 10 random addition sequence replicates was retained and topological robustness (i.e., nodal support) was assessed with rapid bootstrapping with the number of replicates determined by the autoMRE criterion. Concatenation of remaining sequences to assemble the data matrix henceforth referred to as the “original full dataset” was performed using FASconCAT-G (Kück and Longo, 2014).

Because compositional heterogeneity (Delsuc et al., 2005; Jermiin et al., 2004; Kocot et al., 2017; Nesnidal et al., 2010; Rodríguez-Ezpeleta et al., 2007) and branch-length heterogeneity (Kocot et al., 2017; Struck et al., 2014) have been shown to be potential sources of systematic error in phylogenomics, we calculated relative composition frequency variability (RCFV; Zhong et al., 2011) and branch-length heterogeneity score (LB; Struck et al., 2014) for each gene in the original full dataset and assembled data matrices corresponding to the best 50, 100, 200, and 500 genes according to RCFV and LB. This allowed us to examine effects of excluding genes with relatively high compositional heterogeneity or branch-length heterogeneity. Average RCFV was calculated for each gene based on per-taxon RCFV scores calculated in BaCoCa 1.104.r with a subclade definition file that divided the taxa into Ambulacraria (Hemichordata + Echinodermata), Vertebrata, Cephalochordata, and Tunicata. LB was calculated for each gene with TreSpEx 1.1 (Struck, 2014) using RAxML single-gene trees generated as described above. As above, concatenation was performed using FASconCAT-G (Kück and Longo, 2014). For brevity, these matrices are referred to using abbreviated names such as LB_50, which represents the best 50 genes according to branch-length heterogeneity or RCFV_200, which represents the best 200 genes according to RCFV.

Maximum likelihood (ML) analyses were conducted for all data matrices in RAxML 8.2.8 (Stamatakis 2014). Matrices were partitioned by gene and the PROTGAMMAAUTOF model was specified for all partitions. The tree with the best likelihood score after 10 random addition sequence replicates was retained and nodal support was assessed with rapid bootstrapping with the number of replicates determined by the autoMRE criterion.

We also conducted Bayesian inference (BI) analyses in Phylobayes MPI 1.6j (Lartillot et al., 2013) using a site-heterogeneous mixture model. Specifically, the CAT+GTR+r4 model (Lartillot and Philippe, 2004) was used to account for site-specific rate heterogeneity (-cat -gtr - dgam 4). Because of the computationally intensive nature of Phylobayes analyses using this model, BI was only conducted for the RCFV_50 and LB_50 data sets. For the BI analysis of RCFV_50, five parallel chains were run for roughly 15,000-18,000 cycles with the first 10,000 discarded as burn-in. For the BI analysis of LB_50, four parallel chains were run for roughly 17,000-28,000 cycles with the first 10,000 discarded as burn-in. The bpcomp maxdiff values (0.0039 for RCFV_50 and 0.0697 for LB_50) were used to assess convergence of chains.

Because the CAT mixture model implemented in Phylobayes could not be feasibly applied to all datasets, we also conducted ML analyses in IQ-TREE using the posterior mean site frequency (PMSF) model (Wang et al. 2017). PMSF is a rapid approximation of the time- and memory-intensive profile mixture model of Lee et al. (2008), which is a variant of the Phylobayes CAT model. Specifically, the LG+C60+G+F model was specified. Because this approach requires a guide tree to infer the site frequency model, we used the previously generated RAxML tree for each PMSF analysis. For the PMSF analyses of RCFV_50 and LB_50, we additionally tested the effect of using the consensus trees recovered by Phylobayes as the guide tree. Nodal support was assessed with 1000 replicates of ultrafast bootstrapping (bb 1000).

To screen for outlier genes and taxa (genes or taxa that have contamination and/or paralogy issues), single-gene RAxML trees were analyzed in Phylo-MCOA (De Vienne et al., 2012) finding successive decomposition axes from individual ordinations (derived from distance matrices) that maximize a covariance function. For detection of “complete outliers” (genes or taxa most likely to have contamination and/or paralogy issues) we used values of k=1.5 and thres=0.5.

After the above analyses were conducted, transcriptome data became available for two additional tunicate species: *Clavelina lepadiformis* (Aplousobranchia) and *Salpa thompsoni* (Thaliacea). We assembled an additional dataset (Full dataset+2) following an identical approach to that described above to produce the original full dataset except for the addition of these two taxa. The minimum number of taxa required to keep a gene was kept at fifteen (>50% of the original 28 sampled taxa). We also assembled datasets corresponding to the best 50 genes according to RCFV (RCFV_50+2) and LB (LB_50+2) identified as described above that were retained by our pipeline after the addition of these two taxa (47 and 48 genes, respectively). Maximum likelihood analyses were conducted on these datasets in RAxML as described above.

We examined tree certainty (TC), relative tree certainty (RTC), and internode certainty (IC) using the approaches of Salichos and Rokas (2013) and Salichos et al. (2014) as calculated in RAxML 8.2.4. We calculated TC and RTC for the original full dataset and each of the submatrices based on this dataset. Trees resulting from the RAxML analysis of each dataset (provided to RAxML with “-t”) and trees based on the corresponding RAxML single-gene trees (provided with "-z") were used. IC was calculated for the original full dataset based on the corresponding RAxML single-gene trees (provided with "-z"). TC, RTC, and IC were calculated under stochastic bipartition adjustment (excluding conflicting bipartitions).

Finally, we sought to confirm species identification by extracting cytochrome c oxidase subunit I (COI) sequences from all of our newly generated transcriptomes analyzed herein (when present) and conducting a phylogenetic analysis with publicly available tunicate COI sequences. Sequences were manually put in the proper open reading from and were translated to amino acids using the ascidian mitochondrial code in MEGA 7.0.14 (Kumar et al., 2016). Amino acids were aligned with MUSCLE (Edgar, 2004) as implemented in MEGA 7.0.14. A phylogenetic analysis was conducted in RAxML using the approach described above for our concatenated data matrices except the alignment was not partitioned by gene.

## 3. Results

Our bioinformatic pipeline resulted in a matrix (“original full dataset”) of 798 genes totaling 254,865 amino acid (AA) positions in length (Table 4). The average Alicut-trimmed alignment length was 319 AAs with the longest being 1,326 AAs and the shortest being 55 AAs. All genes were sampled for at least 15 taxa but some were sampled for all 28 taxa with an average of 24 taxa sampled per gene. Missing data in the original full dataset was 22.57% (i.e., 77.43% matrix occupancy).

**Table 4.** Length, percent missing data, tree certainty (TC), and relative tree certainty (RTC) for each data matrix analyzed.

| **Data matrix** | **Genes** | **Taxa** | **Length**<br>**(AAs)** | **Missing**<br>**data** | **TC** | **RTC** |
| --- | --- | --- | --- | --- | --- | --- |
| Original full dataset | 798 | 28 | 256,609 | 22.6% | 8.76 | 0.35 |
| RCFV_50 | 50 | 28 | 31,260 | 23.0% | 13.81 | 0.55 |
| RCFV_100 | 100 | 28 | 54,689 | 22.0% | 14.22 | 0.57 |
| RCFV_200 | 200 | 28 | 99,576 | 21.9% | 10.14 | 0.41 |
| RCFV_500 | 500 | 28 | 196,200 | 21.9% | 9.78 | 0.39 |
| LB_50 | 50 | 28 | 15,203 | 24.6% | 13.95 | 0.56 |
| LB_100 | 100 | 28 | 32,776 | 23.4% | 16.14 | 0.65 |
| LB_200 | 200 | 28 | 66,032 | 22.6% | 16.02 | 0.64 |
| LB_500 | 500 | 28 | 166,734 | 22.4% | 11.73 | 0.47 |
| Full dataset+2 | 788 | 30 | 258,910 | 22.6% |  |  |
| RCFV_50+2 | 47 | 30 | 29,831 | 22.6% |  |  |
| LB_50+2 | 48 | 30 | 14,990 | 24.4% |  |  |

We conducted ML analyses on all twelve data sets (Figure 1, Supplementary Figures 1-23) and BI analyses on the two smallest data sets: RCFV_50 and LB_50 (Figure 1). All ML and BI analyses recovered Olfactores (Vertebrata + Tunicata) and Tunicata with maximal support (bootstrap support, bs = 100 and posterior probability, pp = 1.00). Within Tunicata, all analyses recovered the appendicularian *Oikopleura dioica* as the sister taxon to the rest of Tunicata with maximal bootstrap support. Of significance, all ML and BI analyses recovered Thaliacea within “Ascidiacea” with maximal support. Specifically, we consistently recovered Stolidobranchia sister to a clade in which Thaliacea was sister to Phlebobranchia and Aplousobranchia. However, a monophyletic Phlebobranchia was only recovered in the analyses of RCFV_50 (Figure 1, Supplementary Figures 1-3) and LB_100 (Supplementary Figures 9-10). Bootstrap support for Phlebobranchia was weak to moderate in the ML analyses recovering it, but this group was strongly supported by posterior probabilities in the BI analysis of RCFV_50 (pp = 0.99). In the BI analysis of LB_50 and all other ML analyses, phlebobranch tunicates were recovered paraphyletic with respect to Aplousobranchia. BI analysis of LB_50 recovered *Corella* sister to Aplousobranchia with strong support (pp = 0.99) but the ML analysis of this matrix weakly supported this relationship. Most other ML analyses recovered *Ascidia* + *Corella* sister to Aplousobranchia. ML Bootstrap support for relationships among phlebobranchs and Aplousobranchia varied, but as more genes (with increasingly poor average per-taxon RCFV or LB scores) were sampled, support for phlebobranch paraphyly increased.

**Figure 1.**
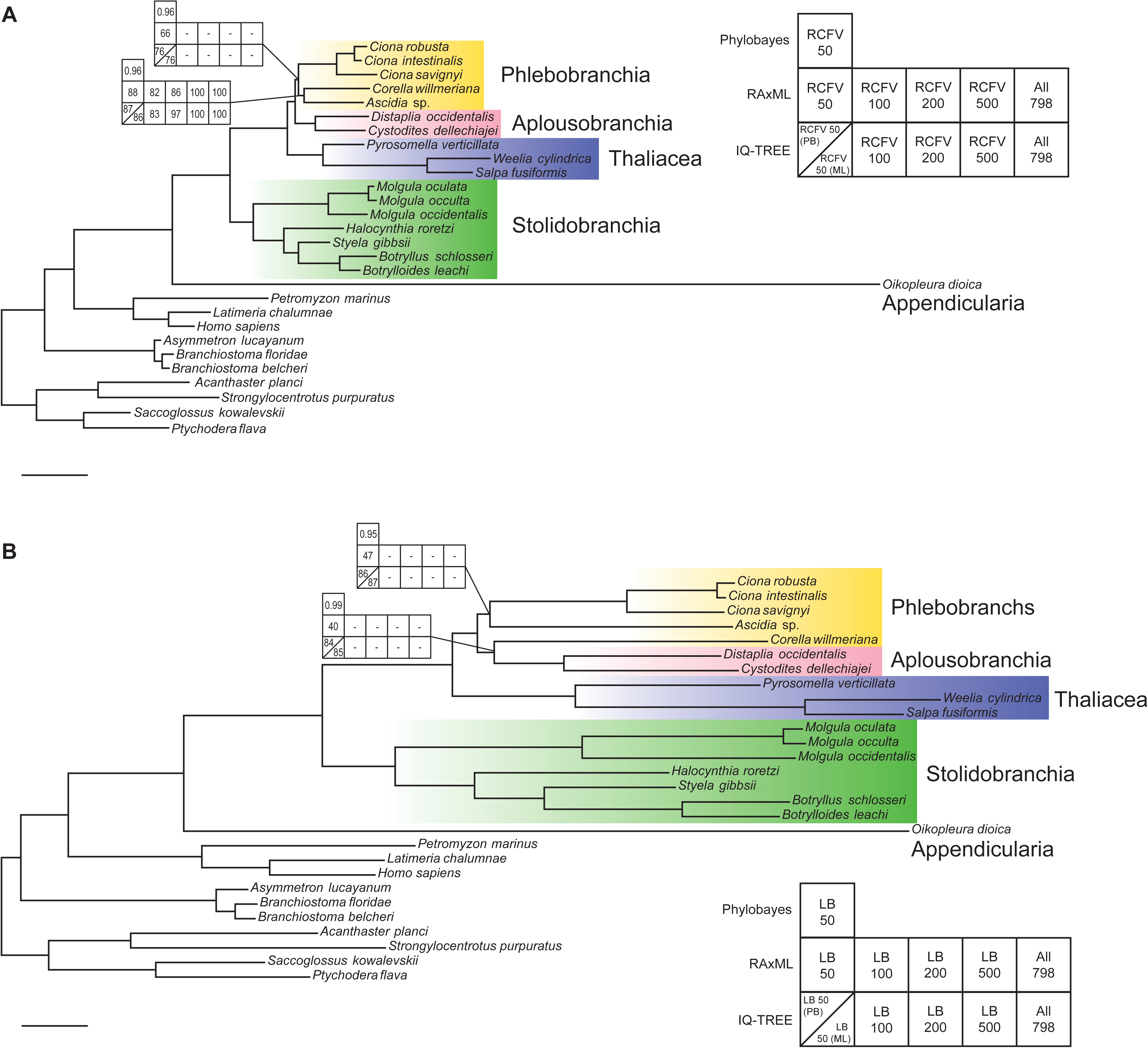
Phylogeny of Tunicata as inferred in the present study. **A.** Consensus phylogram from the Bayesian inference analysis of RCFV_50 with bootstrap support values from RAxML and IQ-TREE ML analyses of RCFV_50, RCFV_100, RCFV_200, RCFV_500, and the original full dataset shown. **B.** Consensus phylogram from the Bayesian inference analysis of LB_50 with bootstrap support values from RAxML and IQ-TREE ML analyses of LB_50, LB_100, LB_200, LB_500, and the original full dataset shown. Nodes without support matrices received maximal support in all BI and ML analyses. For the IQ-TREE analyses of RCFV_50 and LB_50, both the Phylobayes (PB) and RAxML (ML) topologies were tested as guide trees and bootstrap support values resulting from these analyses are presented above and below the diagonal line in the bottom left cells of the support matrices, respectively. Dashes in support matrices indicate that a relationship was not recovered. Scale bars represent 0.1 substitutions per site. Corresponding ML tree topologies are presented in Supplementary Figures 1-22.

Aside from nodes dealing with placement of and relationships among phlebobranch tunicates, all other major tunicate clades as well as relationships within them were consistently recovered and maximally supported in all analyses. Molgulididae was recovered sister to the rest of Stolidobranchia with *Halocynthia* (Pyuridae) sister to Styelidae. Within Styelidae, *Styela* was recovered sister to *Botryllus* + *Botrylloides*. All analyses recovered Thaliacea monophyletic with *Pyrosomella* sister to a clade consisting of the salps Iasis *(=Weelia)* and *Salpa.* Likewise, all analyses recovered Aplousobranchia monophyletic.

ML analyses of a data matrix assembled in the same manner as the original full dataset but with the addition of transcriptome data from *Clavelina lepadiformis* (Aplousobranchia) and *Salpa thompsoni* (Thaliacea; “full dataset+2”; Supplementary Figure 21) resulted in the same general branching order as the analysis of the original full dataset. *C. lepadiformis* was recovered sister to the remaining Aplousobranchia with maximal support, and Aplousobranchia was recovered within Phlebobranchia. *S. thompsoni* was recovered sister to *S. fusiformis* with maximal support. Likewise, analyses of datasets with reduced compositional heterogeneity (RCFV_50+2; Supplementary Figure 22) and branch-length heterogeneity (LB_50+2; Supplementary Figure 23) including these taxa also reflected the topologies recovered in our original analyses without the addition of these data.

ML analyses based on reduced subsets of genes had higher TC and RTC scores than the analysis of the original full dataset (Table 4), which had a TC of 8.76 and an RTC of 0.35. The dataset with the highest tree certainty and relative tree certainty, LB_100, had a TC of 16.14 an RTC of 0.65. Among the smallest two data sets, RCFV_50 (TC = 13.81, RC = 0.55) had slightly lower values than LB_50 (TC = 13.95 and RC = 0.56). We also examined internode certainty (IC) values for the original full dataset (Supplementary Figure 24). IC values were generally low to very low. Notably, the node nesting Aplousobranchia within Phlebobranchia received zero support.

To search for evidence of overlooked paralogy or contamination that could explain the inconsistencies observed among analyses, we screened the original full dataset (without the addition of *Clavelina lepadiformis* or *Salpa thompsoni)* with Phylo-MCOA. This software did not identify any taxa or genes as “complete outliers” (i.e., taxa with contamination or genes with overlooked paralogs) and no individual sequences were identified as outliers (i.e., paralogs; Supplementary Figures 25-26).

Our taxonomic identifications were generally confirmed to at least the genus-level by comparing COI sequences derived from our transcriptomes to publicly available data. Although COI was not recovered in all of our transcriptomes, placement of COI sequences in the tree (Supplementary Figure 27; also see Supplementary Data on Dryad) was consistent with our identifications and the current taxonomy of the sampled taxa in most cases. The only exception was for the Antarctic tunicate we identified as *Ascidia* sp. in the field. The COI sequence derived from from this tunicate clustered with *Phallusia,* which is in the same family as *Ascidia,* but it is unclear if we misidentified the sampled specimen or if there is a taxonomic problem with these genera. Unfortunately the specimen was destroyed in the field to sample internal tissues for molecular work.

## 4. Discussion

### 4.1 Higher-level tunicate phylogeny

Our phylogenomic analyses recovered Appendicularia sister to all other tunicates and confirm that thaliaceans are derived ascidians. Stolidobranchia is monophyletic and sister to a clade that encompasses all other ascidians and thaliaceans. However, monophyly of Phlebobranchia is ambiguous in our analyses. Based on these results, the traditional groupings of higher level tunicate taxa should be revisited as it is not surprising that features such as benthic versus pelagic lifestyle, solitary versus colonial habit, or structure of feeding apparatuses are evolutionarily plastic. Importantly, our results show that “Ascidiacea” is not monophyletic as traditionally defined. Thus, this term should be abandoned as a formal taxonomic name as it represents a paraphyletic group of benthic tunicates, or it should be redefined to also include Thaliacea.

Of particular interest, the taxonomic composition of Phlebobranchia has not been well circumscribed. Cionidae, which includes the widely-studied model species *Ciona intestinalis, Ciona robusta* and *Ciona savignyi,* was originally included within Phlebobranchia (Berrill, 1936), but this family is viewed as a subclade of Aplousobranchia by some taxonomists (e.g., Kott, 1969, 1990). This family was recovered sister to all other Aplousobranchia in an analysis of mitochondrial cytochrome c oxidase subunit I (COI) by Turon and López-Legentil (2004) and biochemical analyses of vanadium oxidation state by Hawkins et al. (1983) also suggest a close relationship of Cionidae to Aplousobranchia. In contrast, none of our analyses recovered Cionidae within or sister to Aplousobranchia, even though Phlebobranchia was recovered paraphyletic with respect to Aplousobranchia in some analyses. Conclusions about the monophyly of Phlebobranchia are difficult to make based on the analyses presented herein as some trees strongly support the paraphyly of this group and others recover it monophyletic. Interestingly, the dataset with the highest TC and RTC, LB_100, was one of just two data sets that resulted in trees recovering Phlebobranchia monophyletic.

One challenge of previous molecular phylogenetic studies of ascidians has been that aplousobranchs show elevated rates of nucleotide substitution in ribosomal and mitochondrial genes when compared to most other tunicates, leading to concerns about long branch artifacts (e.g., Turon and López-Legentil, 2004; Tsagkogeorga et al., 2009). Many species of aplousobranchs have rapidly-evolving 18S genes with large insertions in multiple parts of the molecule when compared to other tunicates. However, in all of our reconstructed trees based on nuclear protein-coding genes, the sampled aplousobranchs have comparable branch lengths to most other tunicates. Notably, species of *Clavelina* and *Distaplia* sampled in the analysis of 18S by Tsagkogeorga et al. (2009) were among the shortest-branched aplousobranch ascidians in that study. However, they also sampled a species of *Cystodites,* which was a rather long-branched taxon in that analysis, suggesting that evolutionary rates of 18S and nuclear protein-coding genes differ in this lineage.

### 4.2 Thaliaceans evolved from a benthic ancestor

Traditionally, tunicates were classified among three classes with the pelagic Thaliacea (pyrosomes, salps, and doliolids) considered a distinct clade from Ascidiacea (e.g., Ruppert et al. 2004, Brusca et al., 2016). All of our results strongly support placement of Thaliacea within the traditional class Ascidiacea as the sister group of Phlebobranchia + Aplousobranchia as recovered by Swalla et al. (2000) and Stach and Turbeville (2002) with 18S rDNA and as hinted at by analyses of 18S by Tsagkogeorga et al. (2009). These results suggest a greater degree of lability in the evolution of benthic versus pelagic lifestyles than traditionally recognized. Given that early cephalochordates, early vertebrates, and larvaceans were swimming organisms (Mallatt and Chen, 2003), a benthic lifestyle must have evolved in the last common ancestor of the ascidian-thaliacean clade. Subsequent to this change in lifestyle, Thaliacea, which is nested well-within a clade of otherwise benthic ascidians, reacquired a pelagic lifestyle, supporting previous assertions that these pelagic tunicates evolved from a benthic ancestor (Swalla et al. 2000).

Historically, Ascidiacea was classified on the basis of the relative position of the gonads with Enterogona including Aplousobranchia and Phlebobranchia, who have gonads closely associated with the gut, and Pleurogona consisting of Stolidobranchia, who generally have gonads distinct from the gut (Garstang, 1928; Perrier, 1898). Thaliacea have gonads associated with the gut, like Aplousobranchia and Phelobobranchia, reinforces the utility of this morphological character that defined Enterogona as noted by Tsagkogeorga et al. (2009).

### 4.3 The phylogenetic position of Appendicularia

Our results are consistent with previous studies recovering Appendicularia outside of “Ascidiacea” (Govindarajan et al., 2011; Stach and Turbeville, 2002; Swalla et al., 2000; Wada, 1998). In studies based on markers such as 18S rDNA (e.g., Govindarajan et al., 2011; Swalla et al., 2000; Tsagkogeorga et al., 2009) these unusual pelagic tunicates tend to have somewhat elevated nucleotide substitution rates and in, phylogenomic analyses (e.g., Delsuc et al., 2006, 2008; this study), *Oikopleura,* the only larvacean from which genomic or transcriptomic data are available, is an extremely long-branched taxon. Thus, recovery of Appendicularia as the sister group of all other tunicates has been questioned as a possible result of long-branch attraction (Swalla et al. 2000). The most recent study examining tunicate phylogeny with the broadest taxon sampling of 18S to date recovered this unusual group within Ascidiacea as the sister group of Stolidobranchia (Tsagkogeorga et al., 2009). However, to further complicate the issue, Appendicularia and Molgulidae tend to have AT-rich 18S sequences, and this shared compositional heterogeneity could be causing an artefactual attraction of these two taxa (Tsagkogeorga et al., 2009).

The branch leading to *Oikopleura* was noticeably longer from root to tip than all other taxa in our analyses of all genes retained by our pipeline and data sets with reduced compositional heterogeneity. However, as data sets with fewer but ‘better’ genes according to branch-length heterogeneity were analyzed, the branch leading to *Oikopleura* decreased in length relative to other sampled taxa, and was even shorter than the branches leading to *Salpa* and *Iasis* in both the ML and BI analyses of LB_50. Given our consistent recovery of Appendicularia as sister to the rest of Tunicata with maximal support even when compositional heterogeneity and long-branch attraction are reduced, we consider Appendicularia to be an early-branching tunicate lineage and not a derived ascidian clade as previously hypothesized.

### 4.4 Future directions

This study represents a first step towards resolving tunicate phylogeny using genomic data, but greatly improved taxon sampling will be needed to begin to gain a full picture of the evolutionary history of this diverse and important group. For example, Oikopleuridae is one of three families of Appendicularia but the only family included in any molecular phylogenetic investigation to date. Likewise, Sorberacea is an enigmatic clade of benthic, deep-sea tunicates that has been considered to be a separate class from other tunicates (Monniot et al., 1975). Whether this clade is indeed a distinct lineage of Tunicata or yet another derived ascidian lineage has never been tested with any source of molecular data, let alone phylogenomics. Moreover, many questions about tunicate evolutionary history at family level have been challenging to address, particularly within Aplousobranchia, which exhibits extreme branch-length heterogeneity for 18S rDNA. As most tunicates in the present study exhibit more-or-less comparable branch lengths, especially when steps are taken to exclude genes with exceptional branch-length heterogeneity scores, phylogenomics appears well-suited to address these evolutionary questions.

## Acknowledgements

This study was supported by University of Alabama start-up funds to KMK. We thank Alison Sweeney and James Townsend for facilitating collection of specimens of Thaliacea and the faculty and staff of Friday Harbor Labs for facilitating collection of Pacific tunicates. The crew of the *R/V Nathaniel B. Palmer* and *R/V Lawrence M. Gould* facilitated collection of Antarctic tunicates through NSF funds to KMH (ANT-1043745 and OPP-0132032). This study is based in part upon work by BJS supported by the National Science Foundation under Cooperative Agreement No. DBI-0939454. Any opinions, findings, and conclusions or recommendations expressed in this material are those of the author(s) and do not necessarily reflect the views of the National Science Foundation. We thank Justin Cox and Emily Pabst for help with RNA extractions. We thank Deb Crocker and Robert Griffin for assistance with the University of Alabama High Performance Computing (UAHPC) system. This is Auburn University Marine Biology Program contribution #169 and Molette Lab contribution #73.

---

**Supplementary Figure 1.**
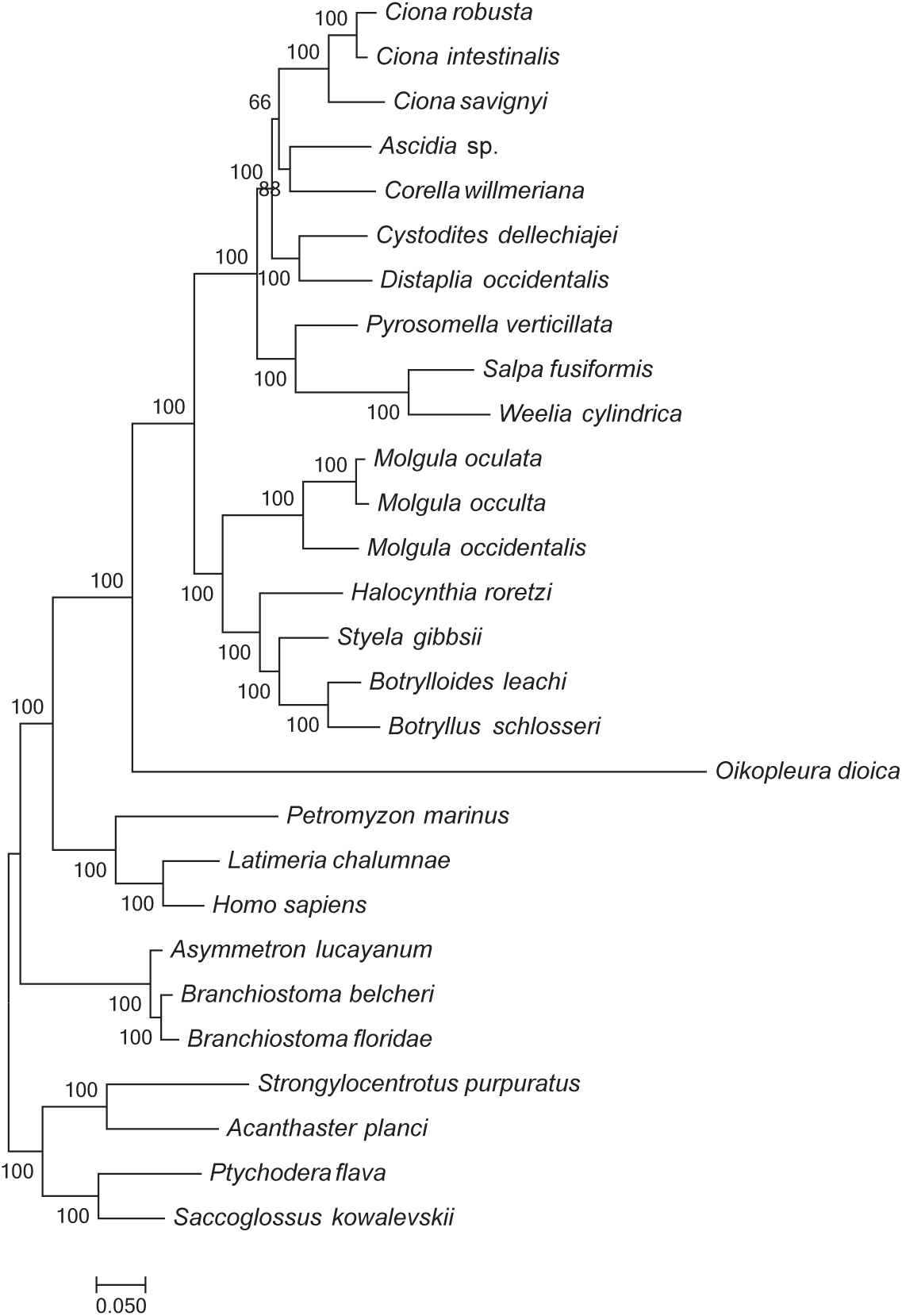
Phylogeny of Tunicata based on the RAxML analysis of the best 50 genes according to RCFV. Bootstrap support values are presented at each node. Scale bar represents 0.1 substitutions per site.

**Supplementary Figure 2.**
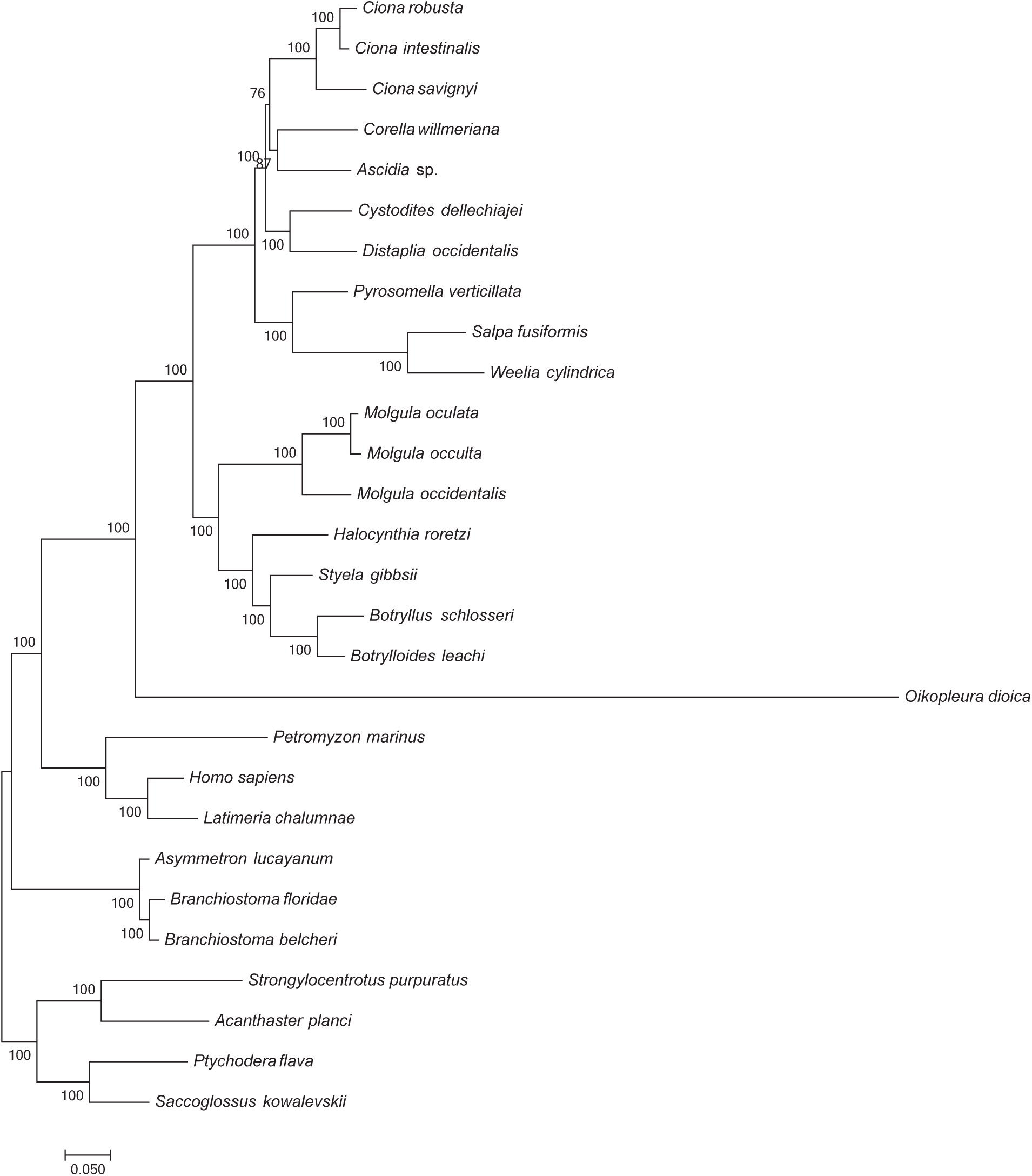
Phylogeny of Tunicata based on the IQ-TREE analysis of the best 50 genes according to RCFV with the RAxML tree used as the guide tree. Bootstrap support values are presented at each node. Scale bar represents 0.1 substitutions per site.

**Supplementary Figure 3.**
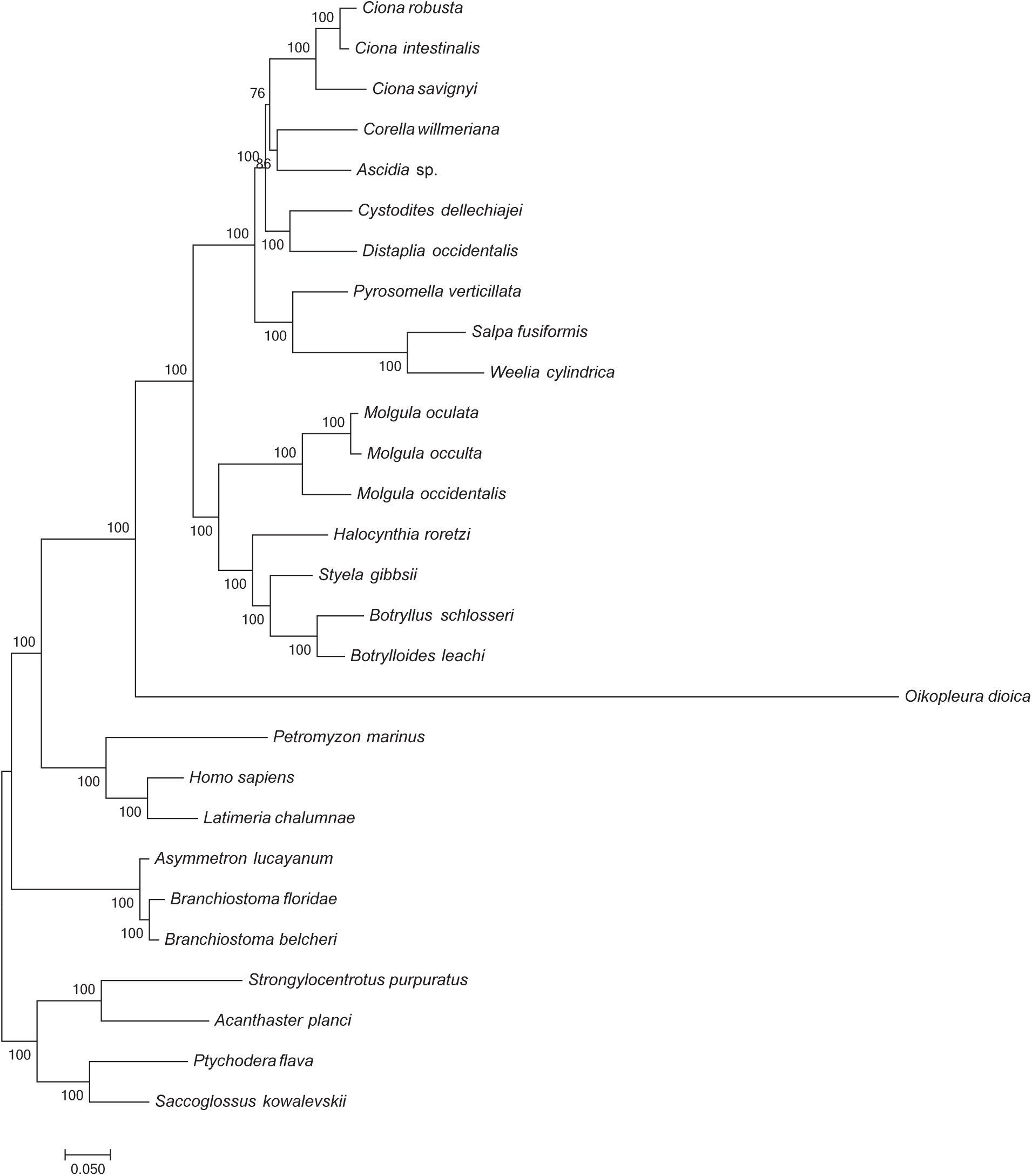
Phylogeny of Tunicata based on the IQ-TREE analysis of the best 50 genes according to RCFV with the Phylobayes tree used as the guide tree. Bootstrap support values are presented at each node. Scale bar represents 0.1 substitutions per site.

**Supplementary Figure 4.**
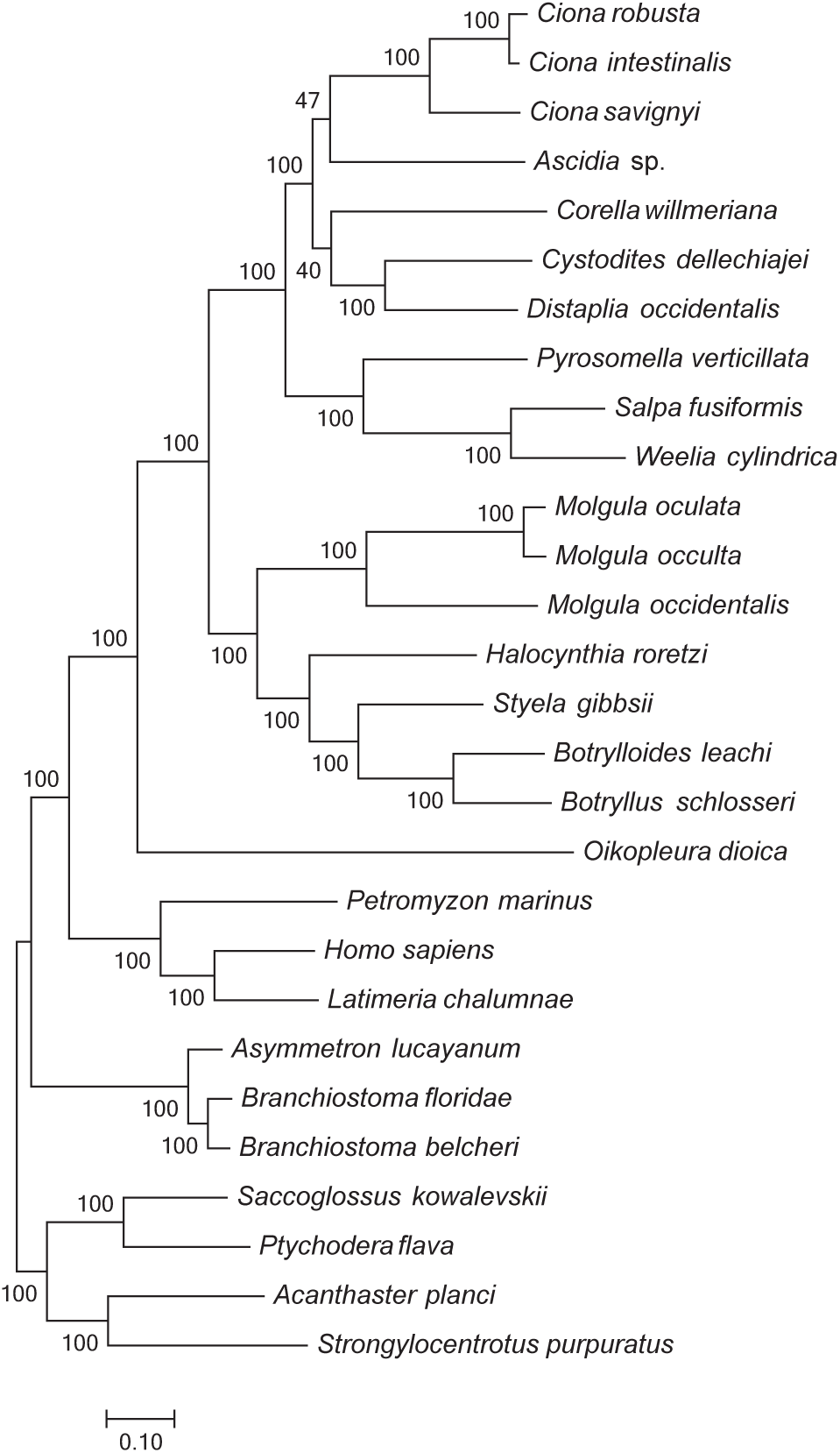
Phylogeny of Tunicata based on the RAxML analysis of the best 50 genes according to LB. Bootstrap support values are presented at each node. Scale bar represents 0.1 substitutions per site.

**Supplementary Figure 5.**
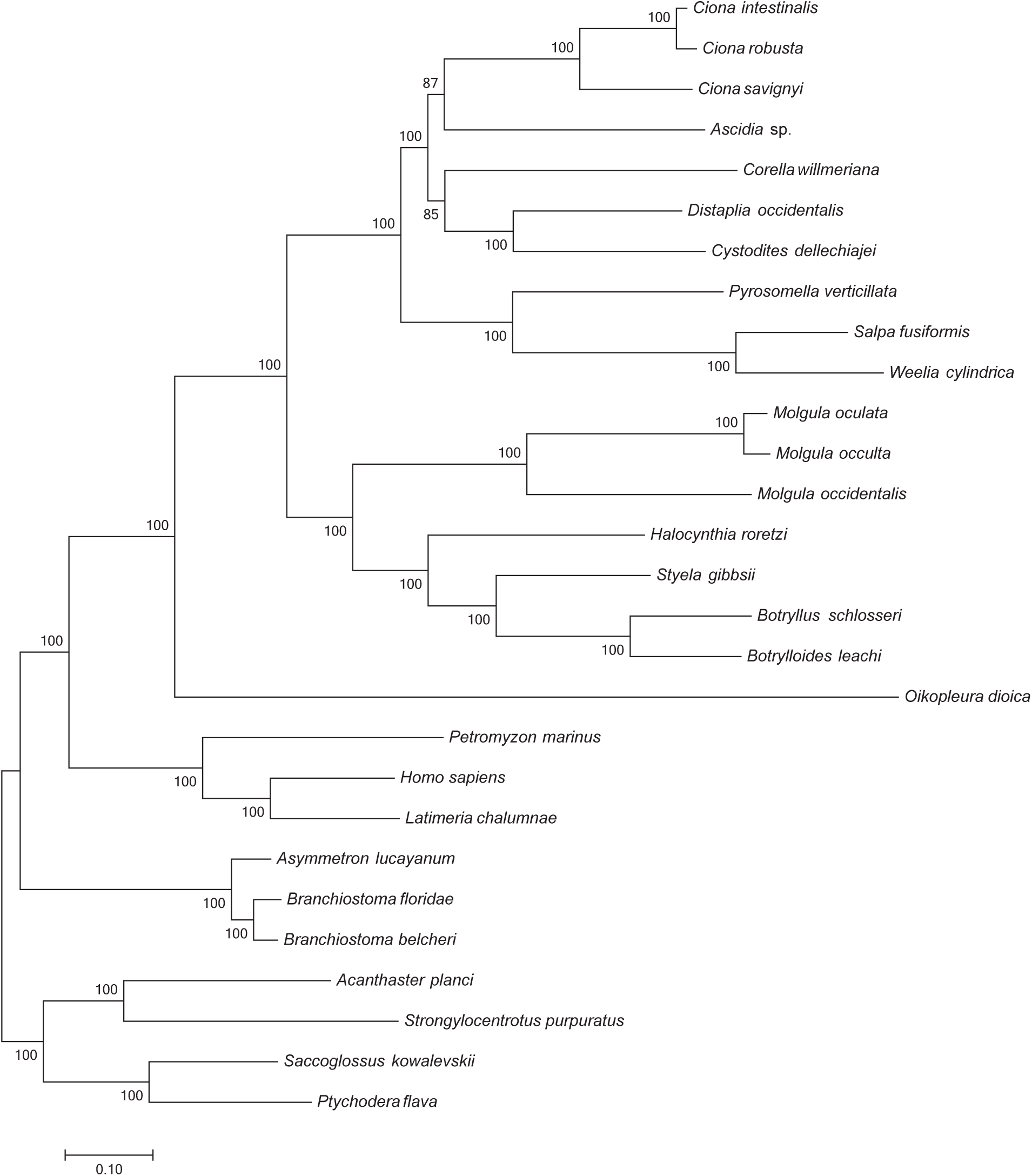
Phylogeny of Tunicata based on the IQ-TREE analysis of the best 50 genes according to LB with the RAxML tree used as the guide tree. Bootstrap support values are presented at each node. Scale bar represents 0.1 substitutions per site.

**Supplementary Figure 6.**
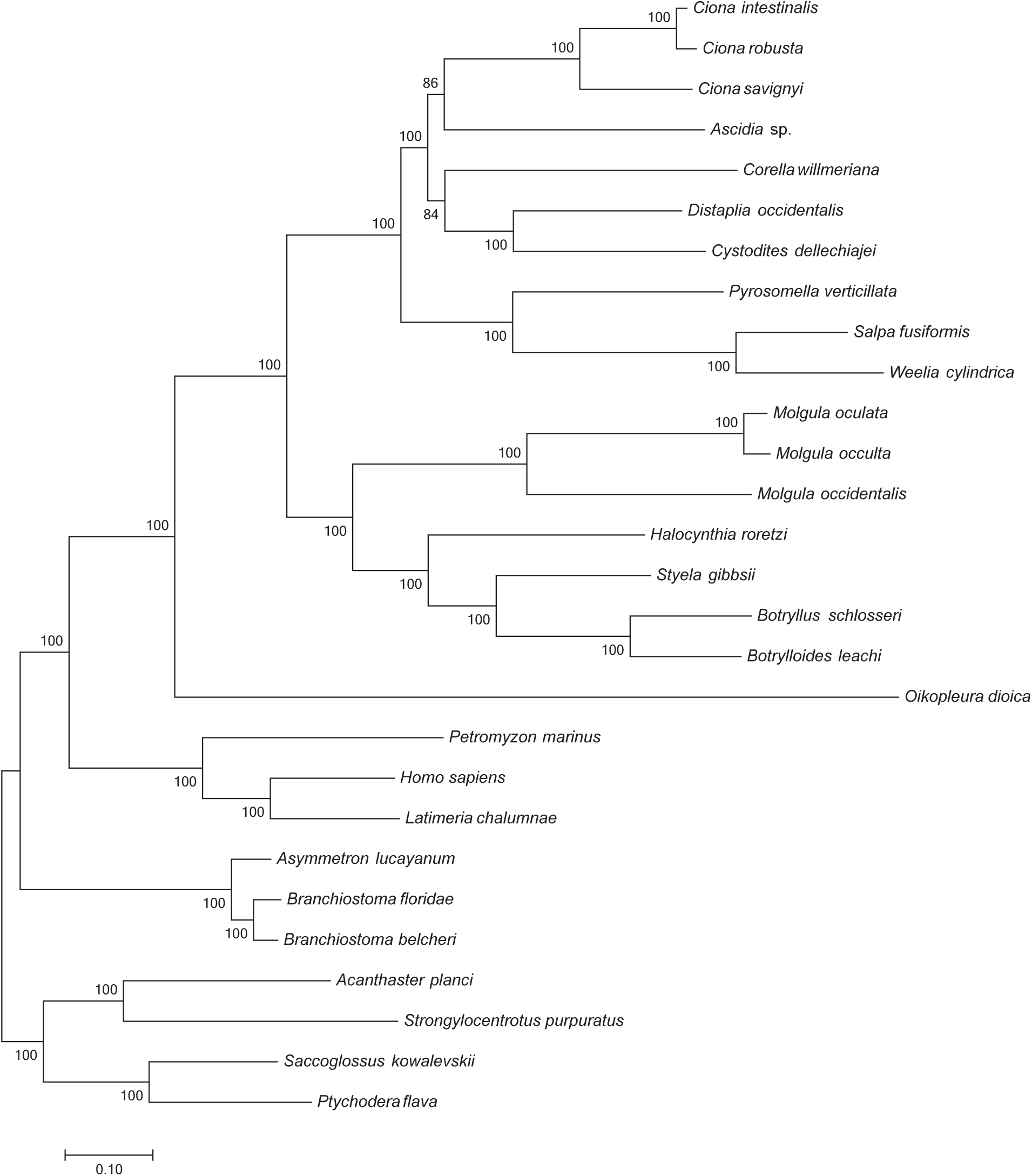
Phylogeny of Tunicata based on the IQ-TREE analysis of the best 50 genes according to LB with the Phylobayes tree used as the guide tree. Bootstrap support values are presented at each node. Scale bar represents 0.1 substitutions per site.

**Supplementary Figure 7.**
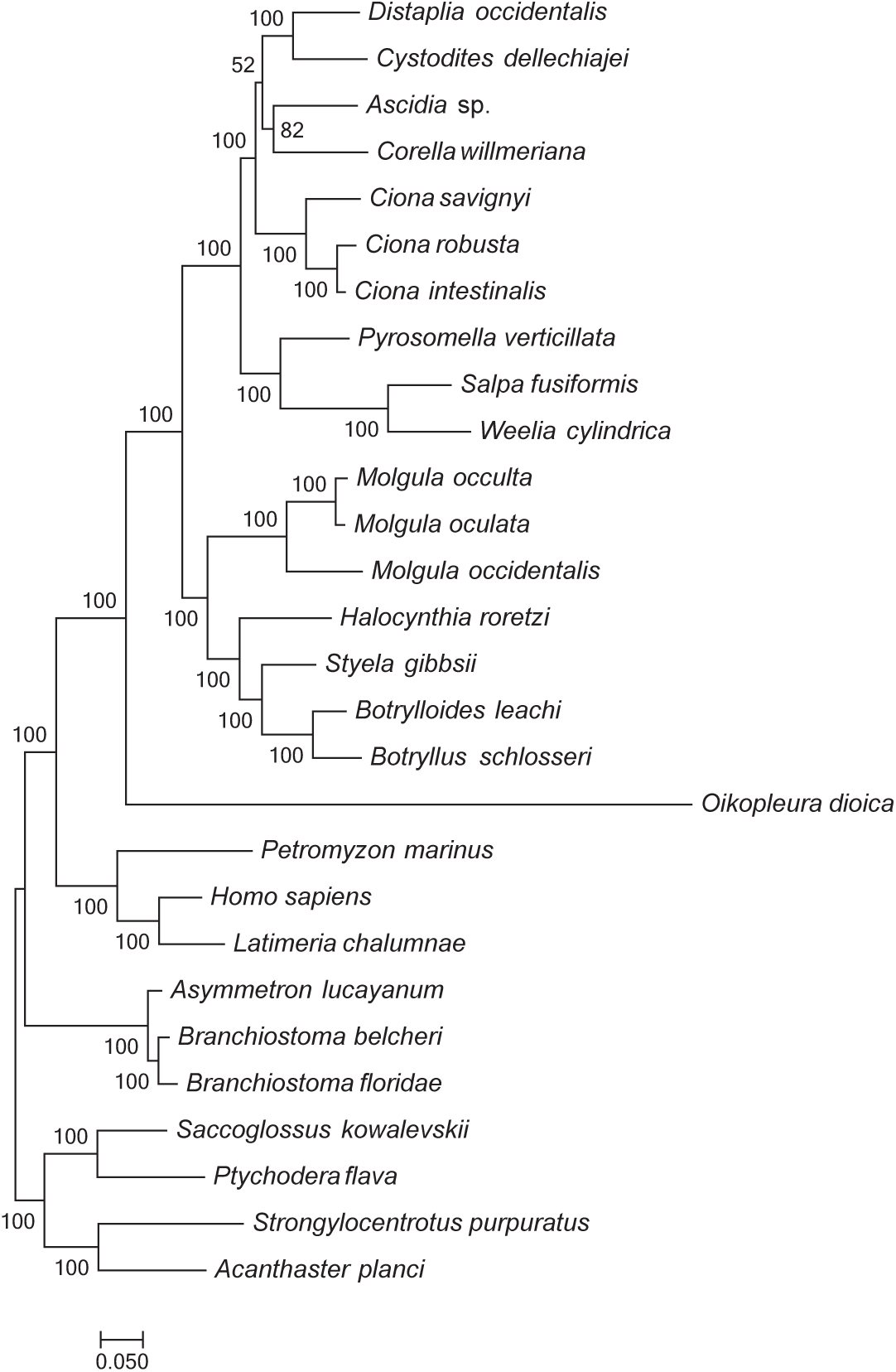
Phylogeny of Tunicata based on the RAxML analysis of the best 100 genes according to RCFV. Bootstrap support values are presented at each node. Scale bar represents 0.1 substitutions per site.

**Supplementary Figure 8.**
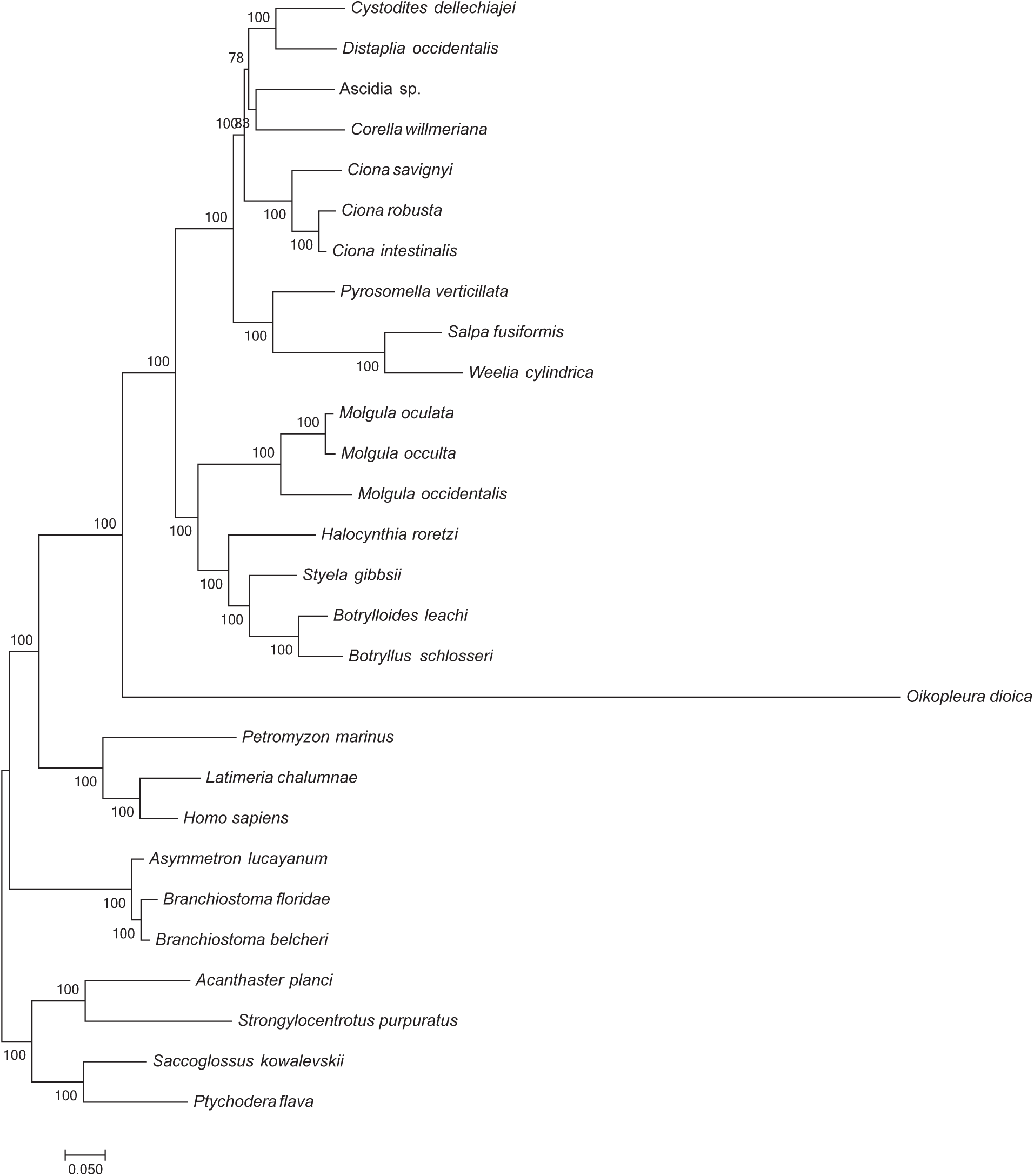
Phylogeny of Tunicata based on the IQ-TREE analysis of the best 100 genes according to RCFV. Bootstrap support values are presented at each node. Scale bar represents 0.1 substitutions per site.

**Supplementary Figure 9.**
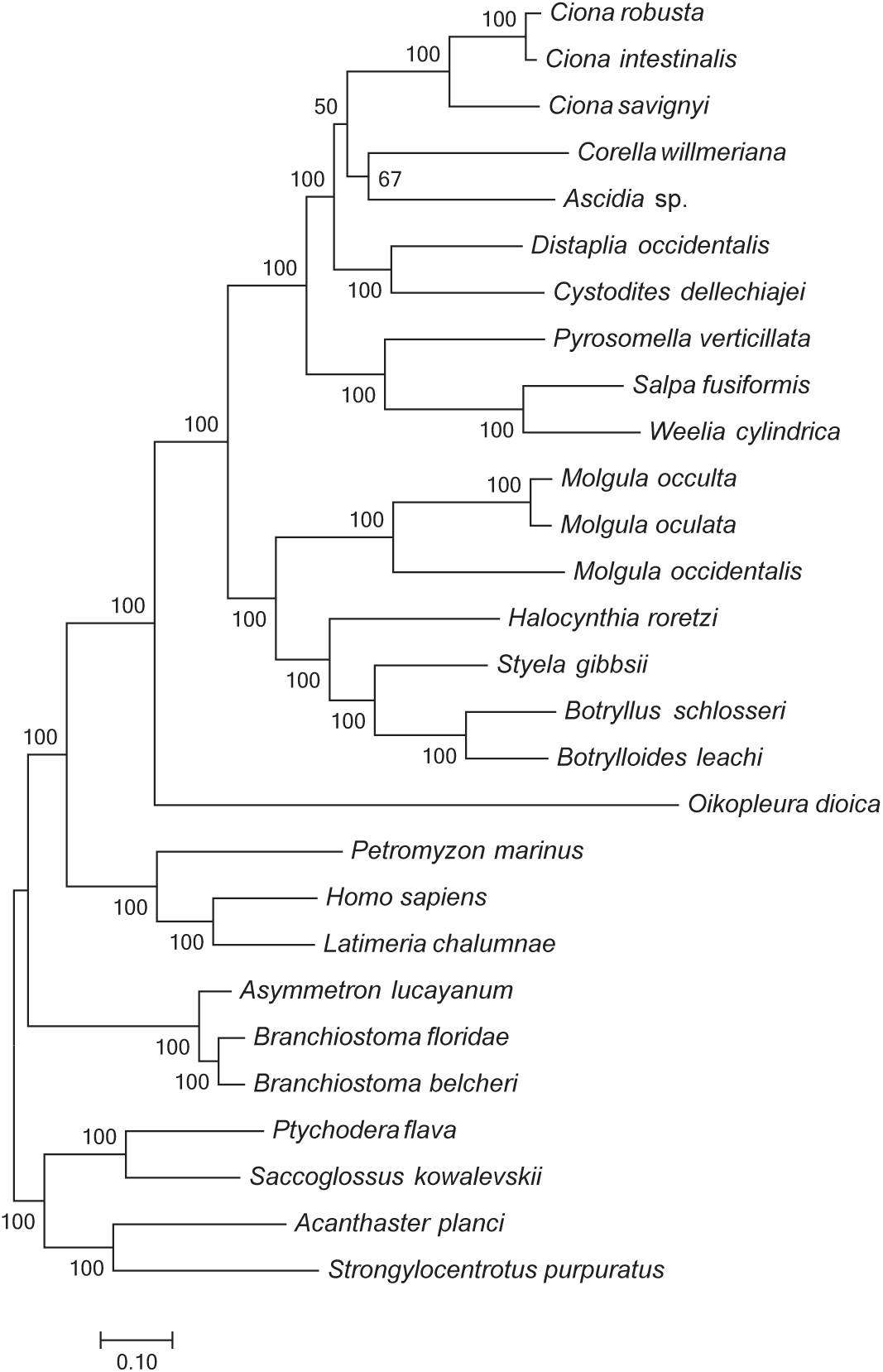
Phylogeny of Tunicata based on the RAxML analysis of the best 100 genes according to LB. Bootstrap support values are presented at each node. Scale bar represents 0.1 substitutions per site.

**Supplementary Figure 10.**
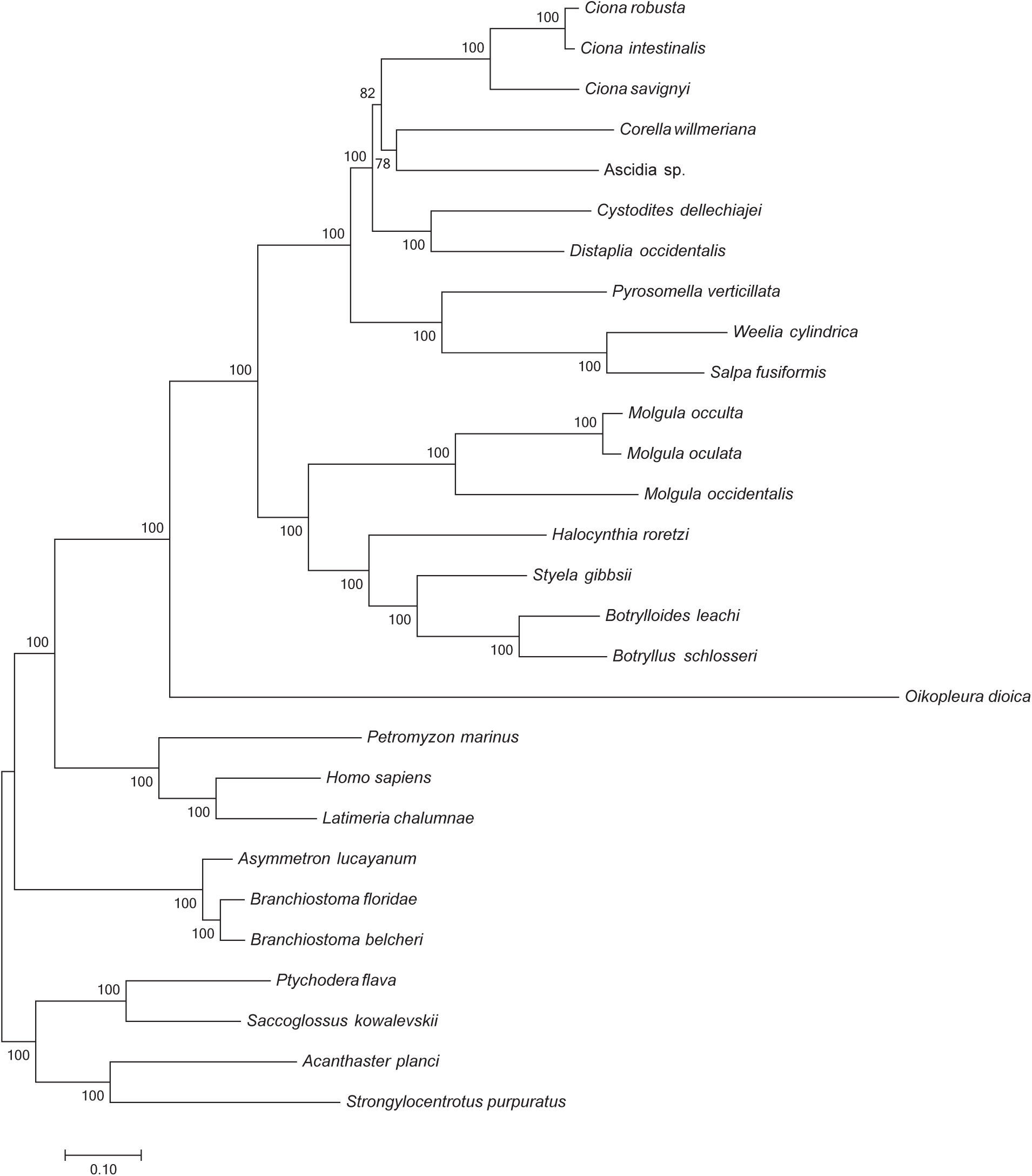
Phylogeny of Tunicata based on the IQ-TREE analysis of the best 100 genes according to LB. Bootstrap support values are presented at each node. Scale bar represents 0.1 substitutions per site.

**Supplementary Figure 11.**
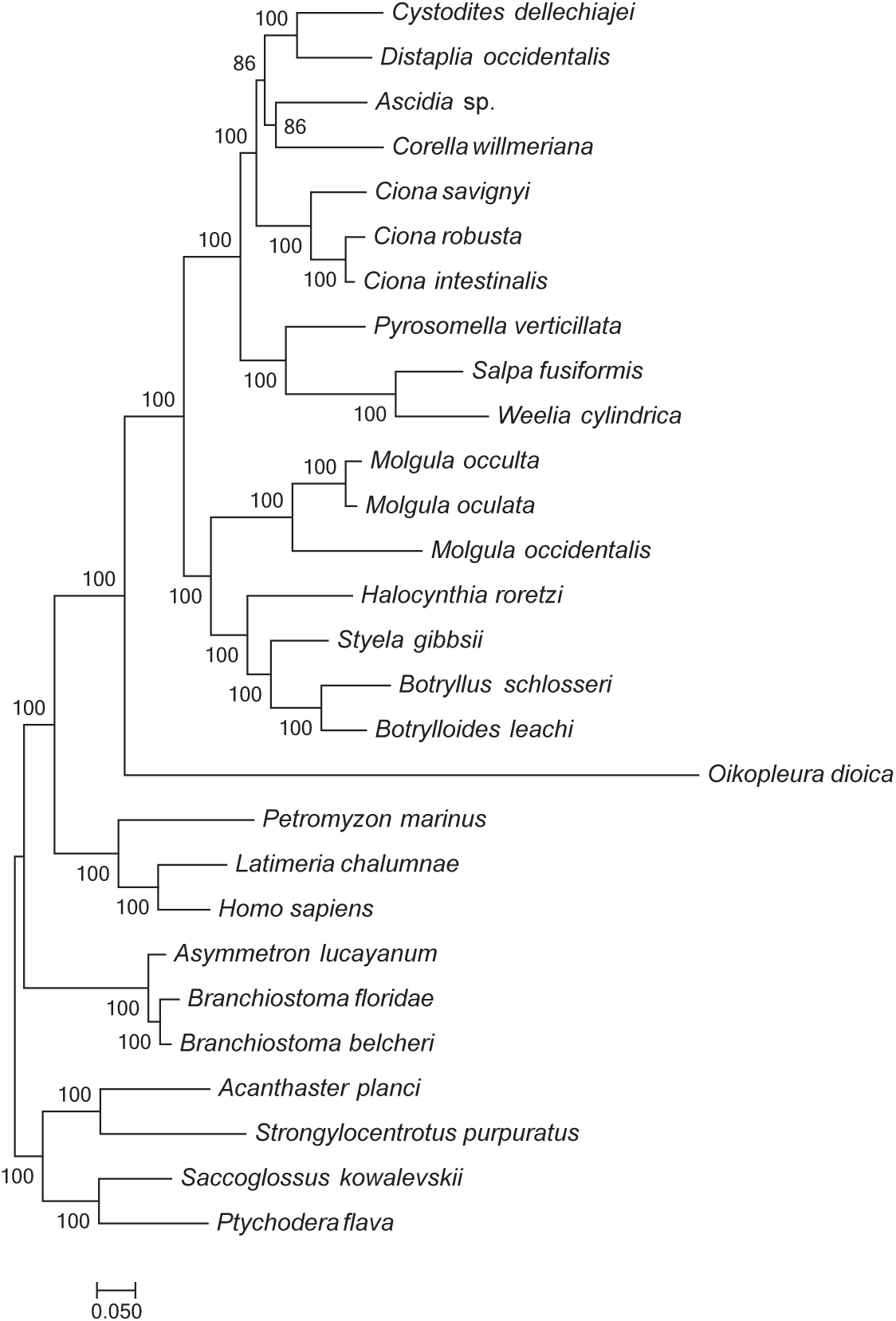
Phylogeny of Tunicata based on the RAxML analysis of the best 200 genes according to RCFV. Bootstrap support values are presented at each node. Scale bar represents 0.1 substitutions per site.

**Supplementary Figure 12.**
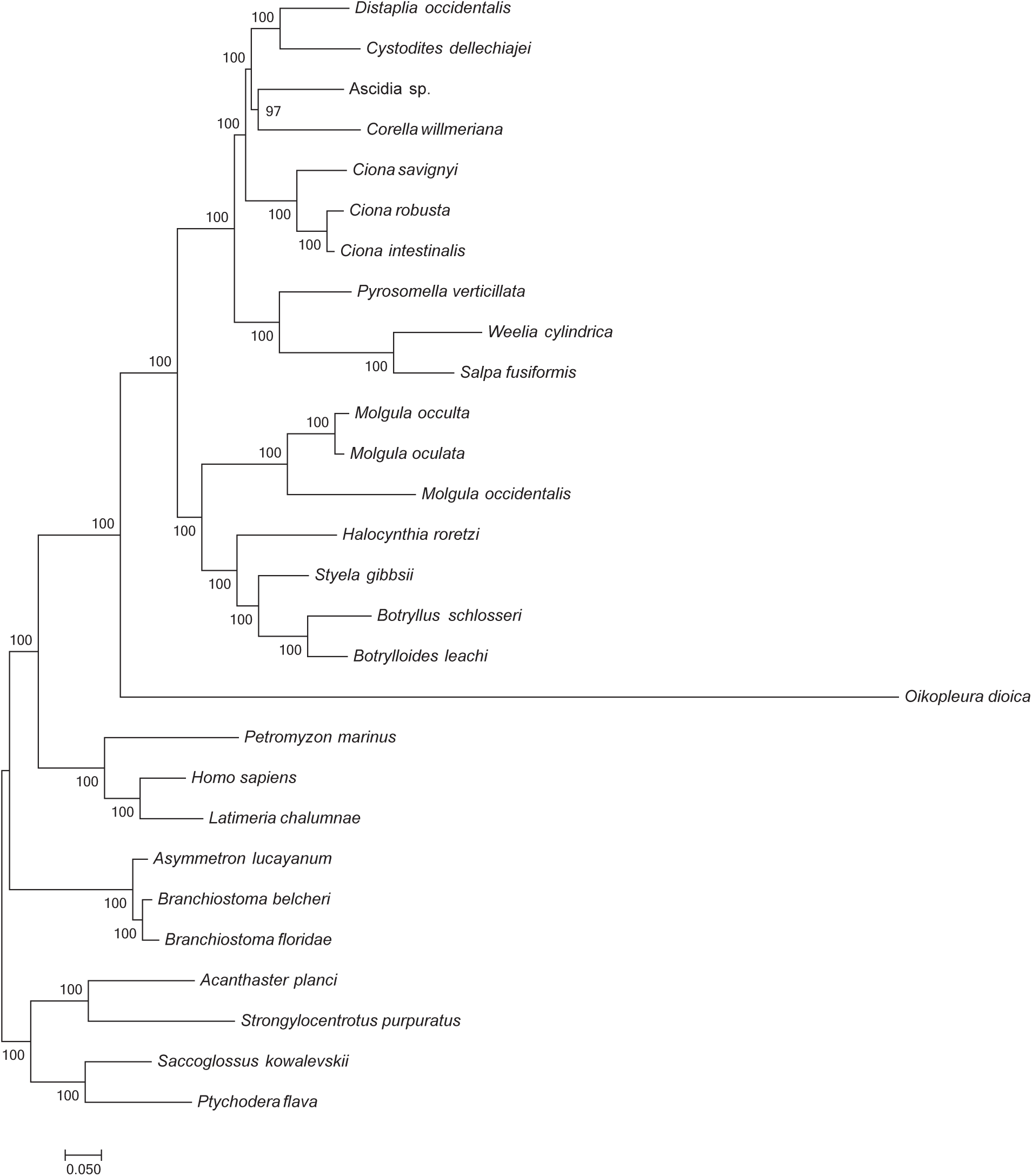
Phylogeny of Tunicata based on the IQ-TREE analysis of the best 200 genes according to RCFV. Bootstrap support values are presented at each node. Scale bar represents 0.1 substitutions per site.

**Supplementary Figure 13.**
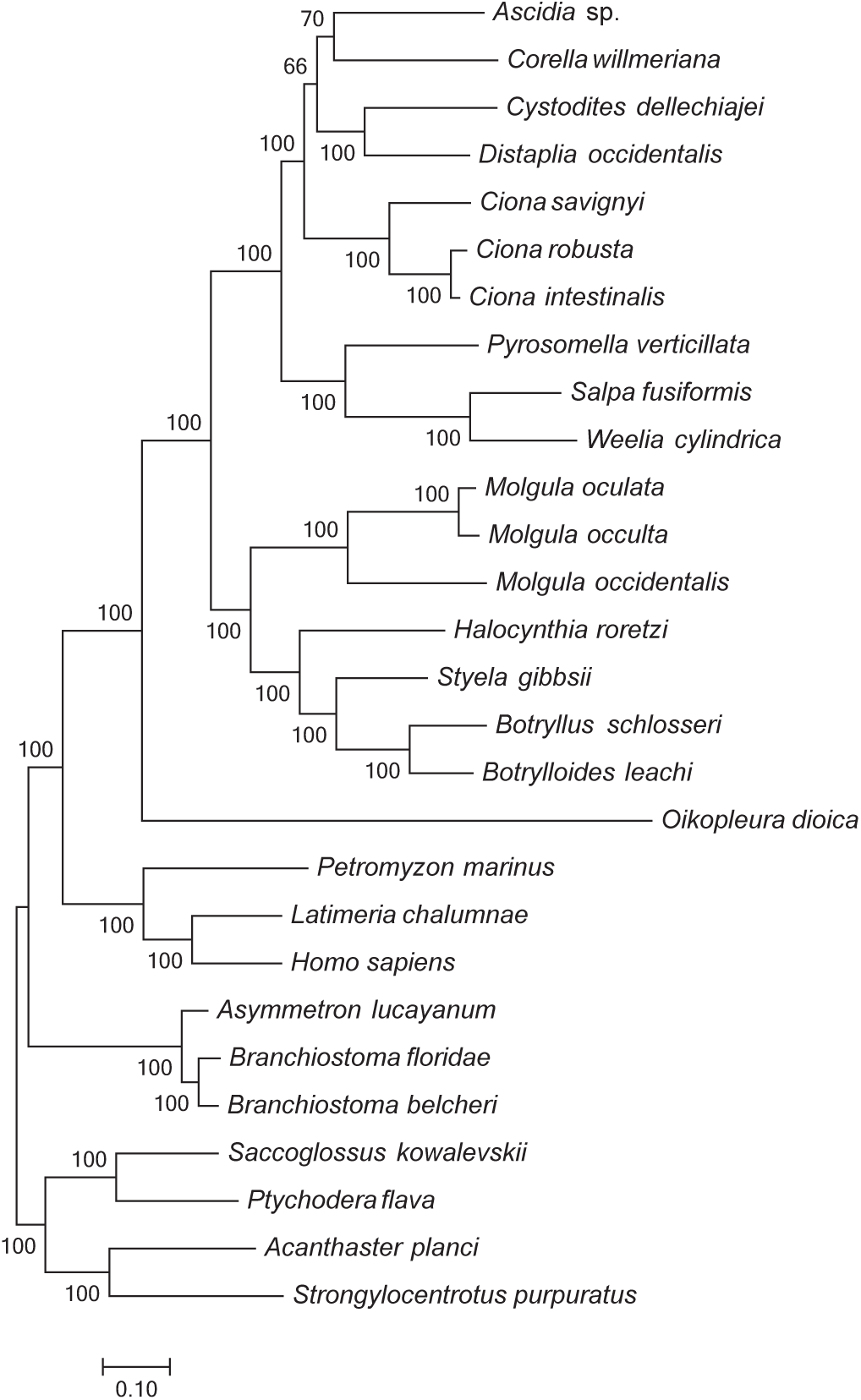
Phylogeny of Tunicata based on the RAxML analysis of the best 200 genes according to LB. Bootstrap support values are presented at each node. Scale bar represents 0.1 substitutions per site.

**Supplementary Figure 14.**
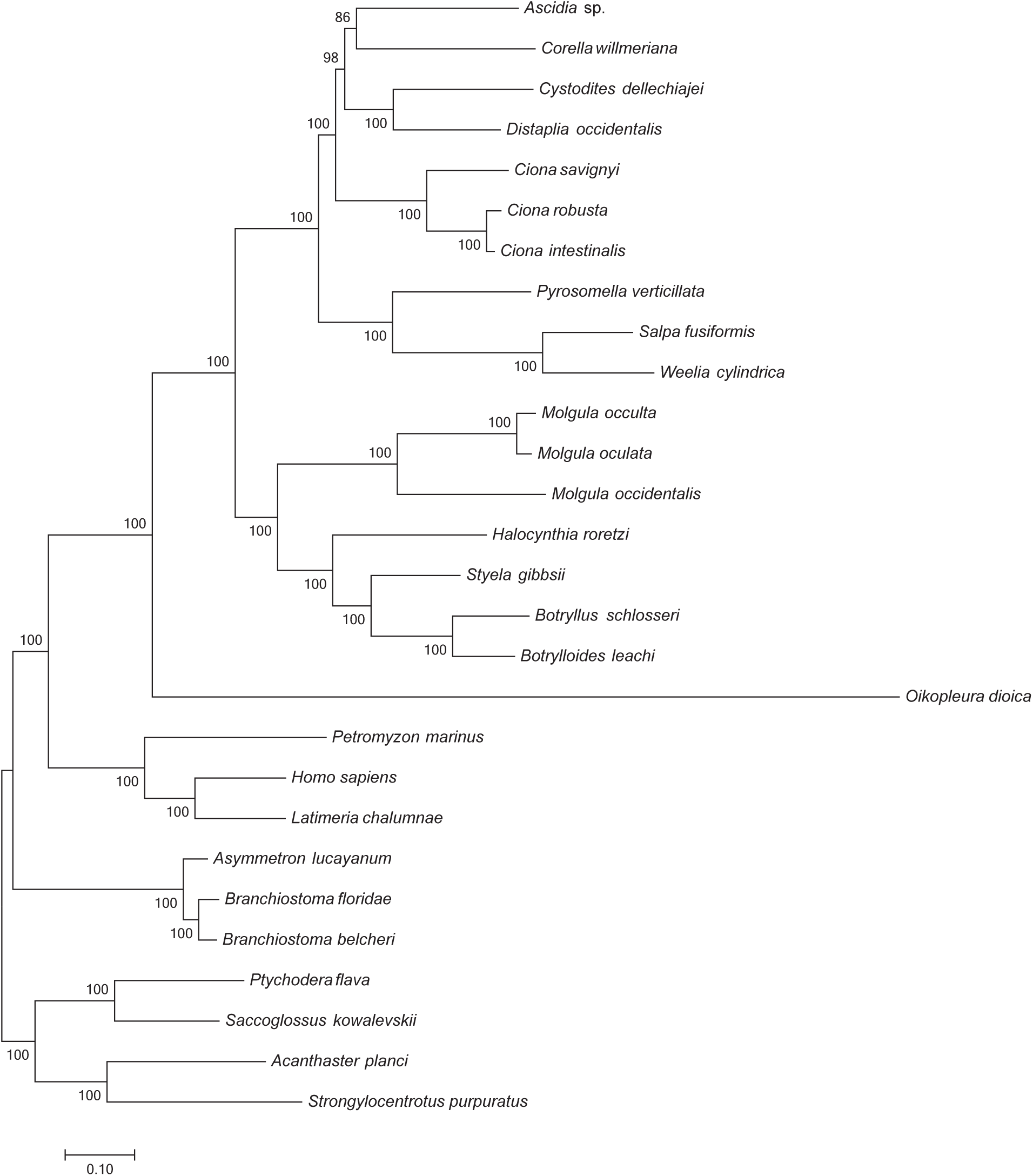
Phylogeny of Tunicata based on the IQ-TREE analysis of the best 200 genes according to LB. Bootstrap support values are presented at each node. Scale bar represents 0.1 substitutions per site.

**Supplementary Figure 15.**
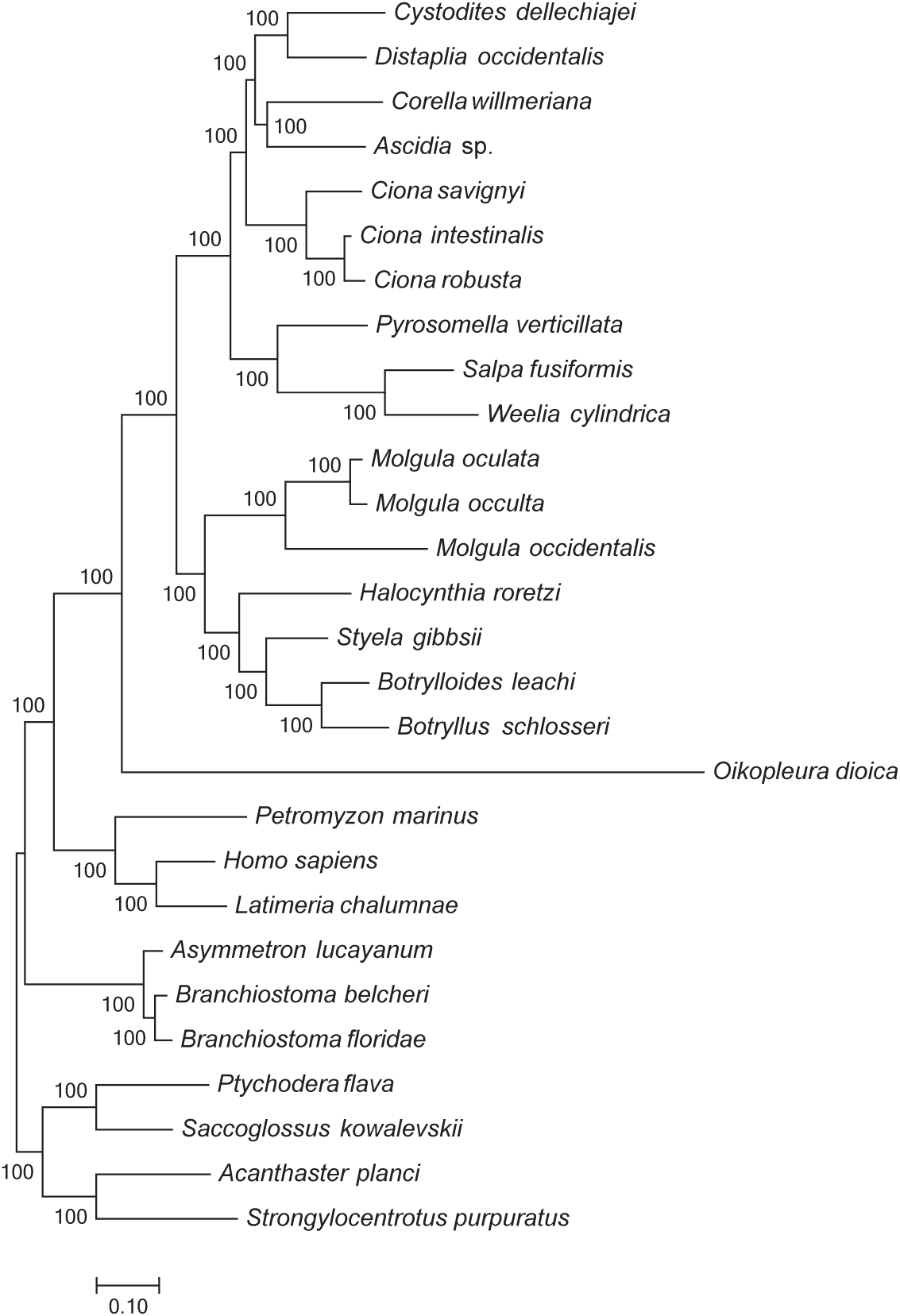
Phylogeny of Tunicata based on the RAxML analysis of the best 500 genes according to RCFV. Bootstrap support values are presented at each node. Scale bar represents 0.1 substitutions per site.

**Supplementary Figure 16.**
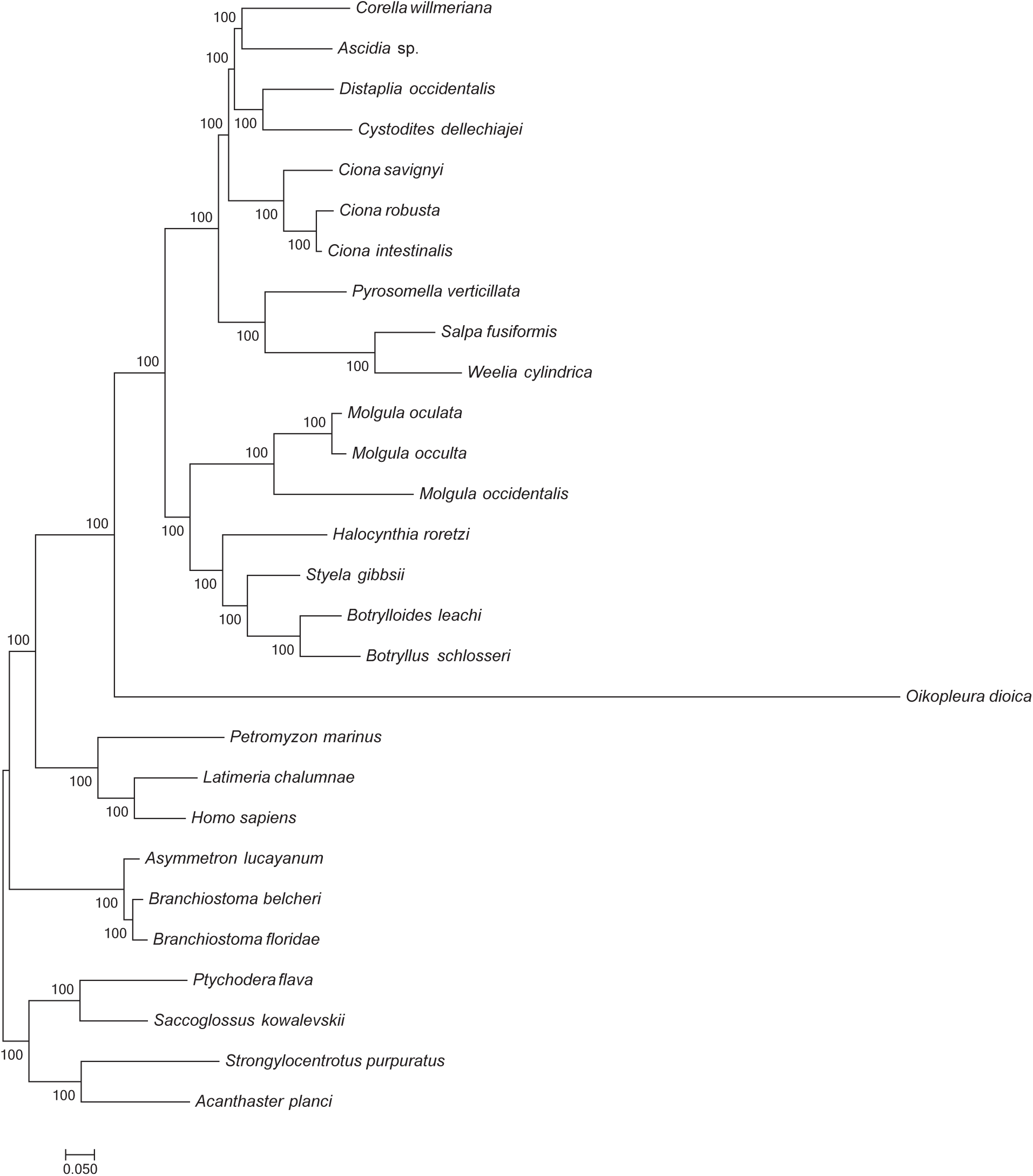
Phylogeny of Tunicata based on the IQ-TREE analysis of the best 500 genes according to RCFV. Bootstrap support values are presented at each node. Scale bar represents 0.1 substitutions per site.

**Supplementary Figure 17.**
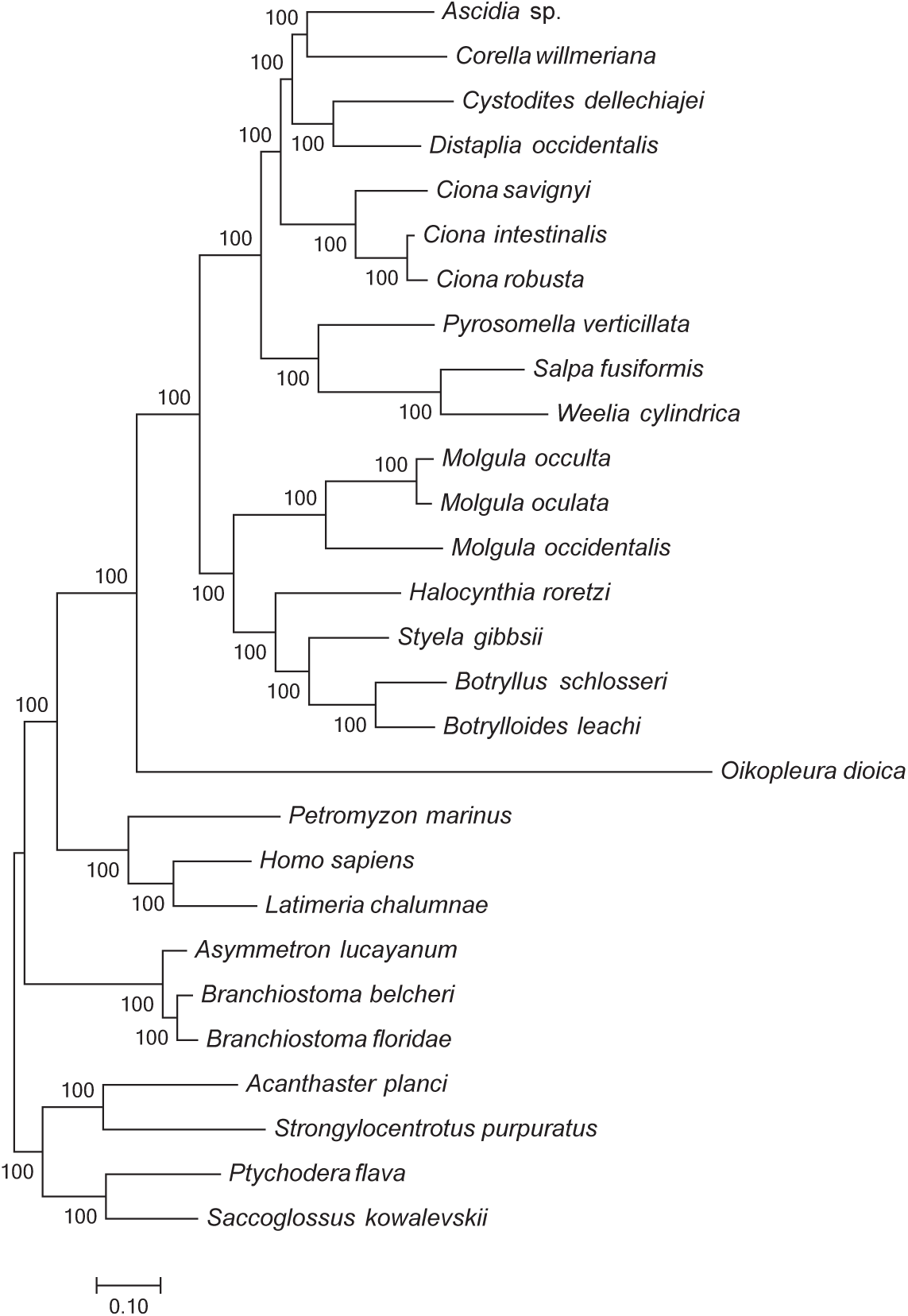
Phylogeny of Tunicata based on the RAxML analysis of the best 500 genes according to LB. Bootstrap support values are presented at each node. Scale bar represents 0.1 substitutions per site.

**Supplementary Figure 18.**
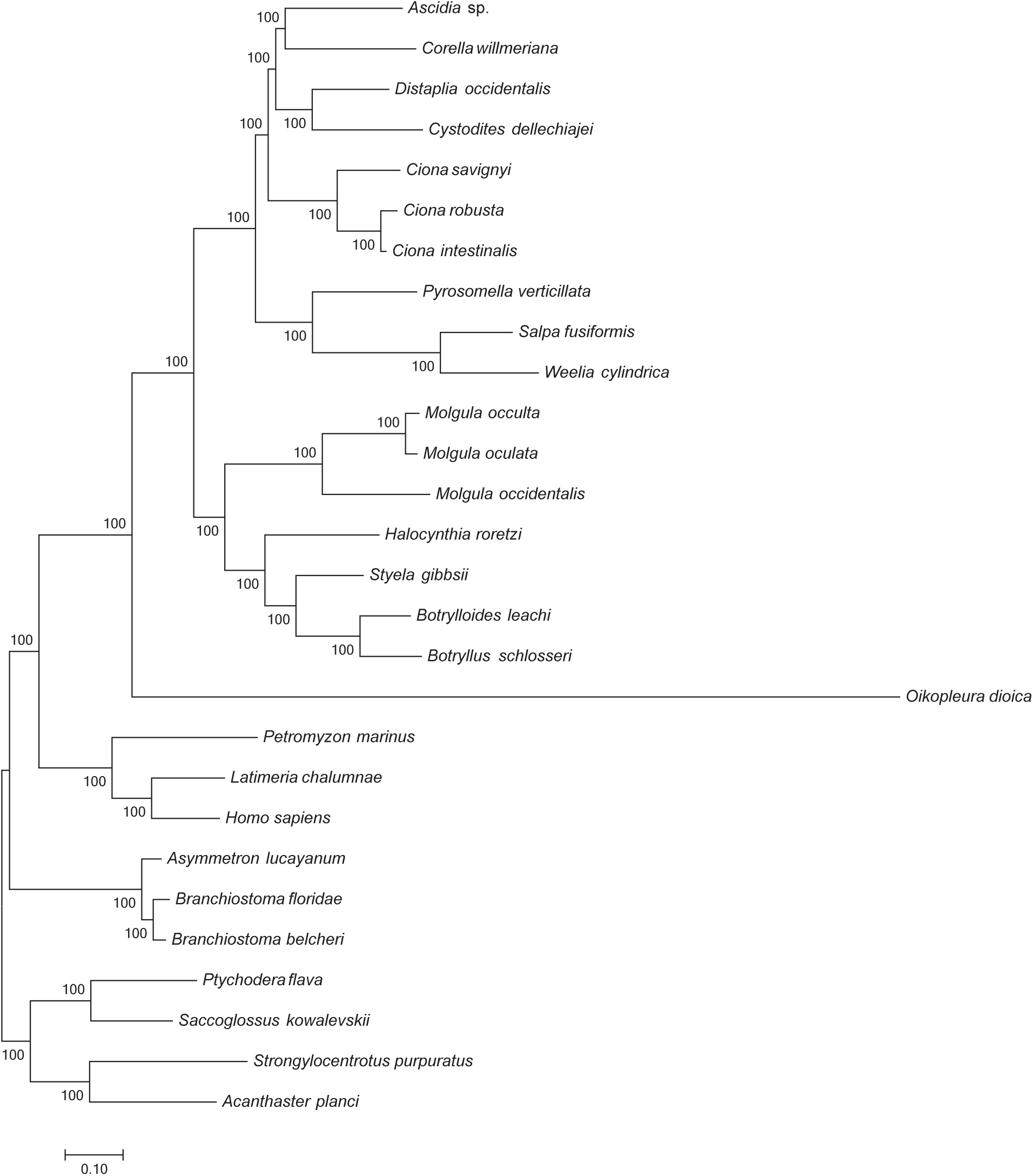
Phylogeny of Tunicata based on the IQ-TREE analysis of the best 500 genes according to LB. Bootstrap support values are presented at each node. Scale bar represents 0.1 substitutions per site.

**Supplementary Figure 19.**
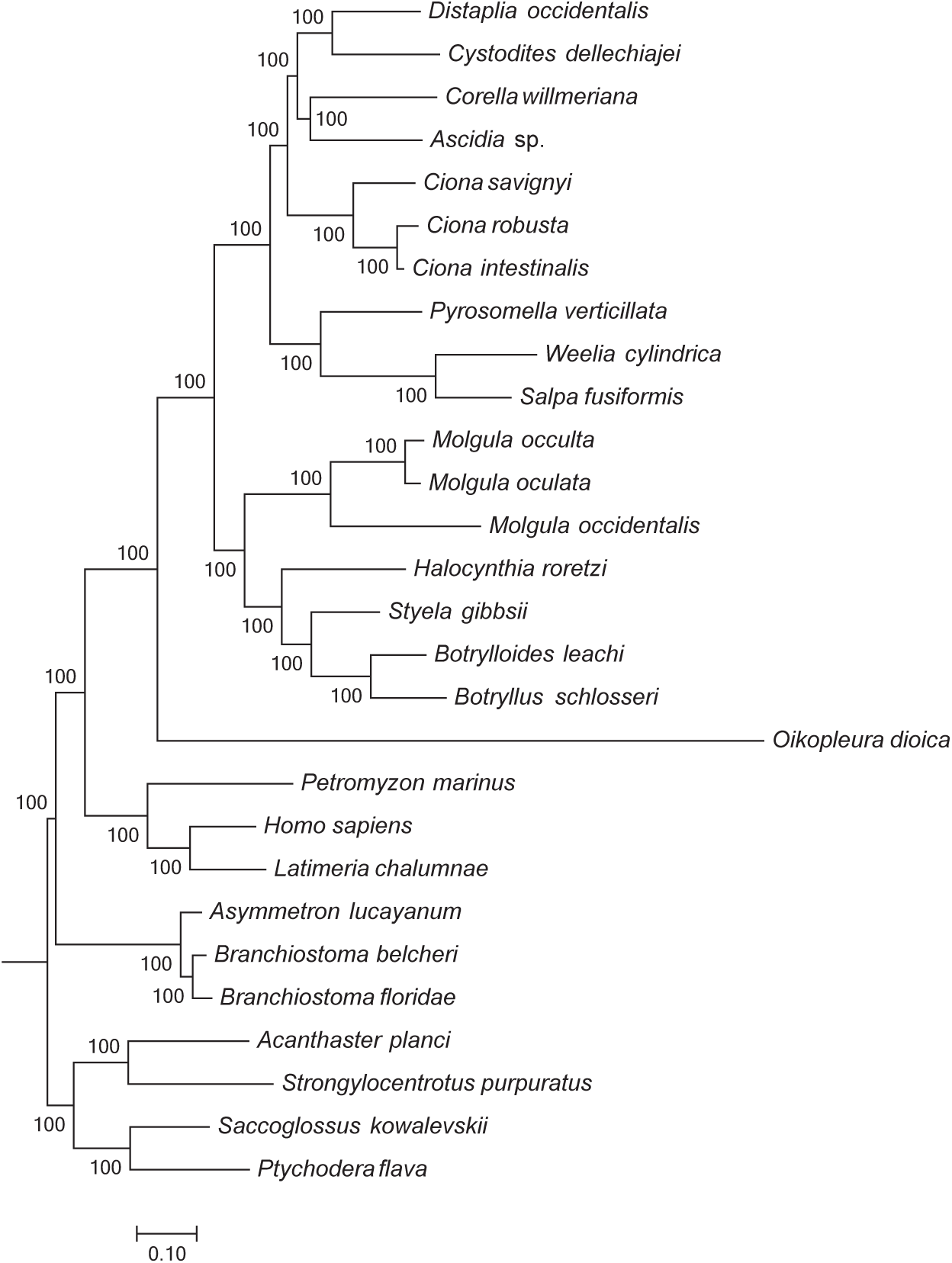
Phylogeny of Tunicata based on the RAxML analysis of the original full dataset (798 genes). Bootstrap support values are presented at each node. Scale bar represents 0.1 substitutions per site.

**Supplementary Figure 20.**
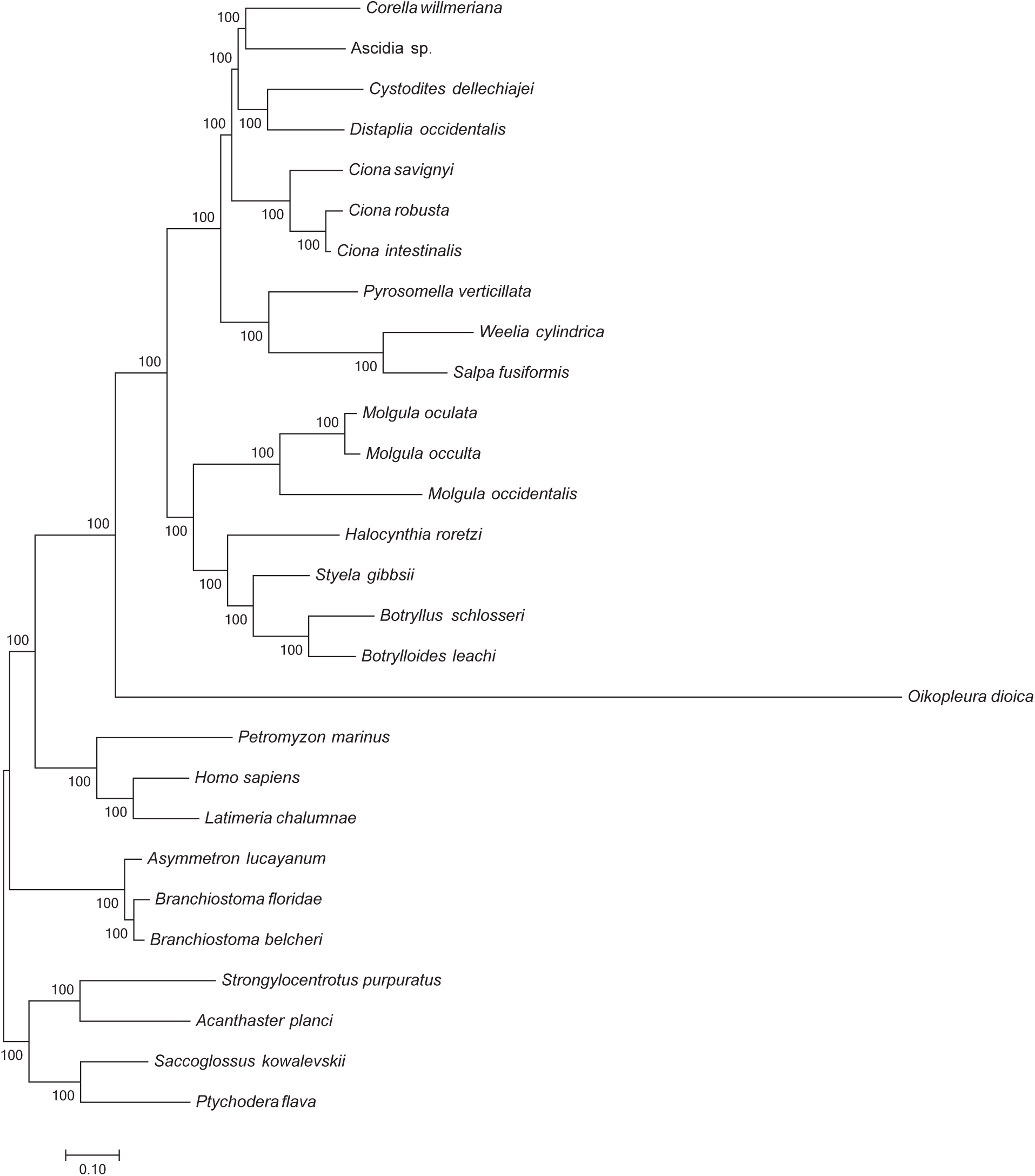
Phylogeny of Tunicata based on the IQ-TREE analysis of the original full dataset (798 genes). Bootstrap support values are presented at each node. Scale bar represents 0.1 substitutions per site.

**Supplementary Figure 21.**
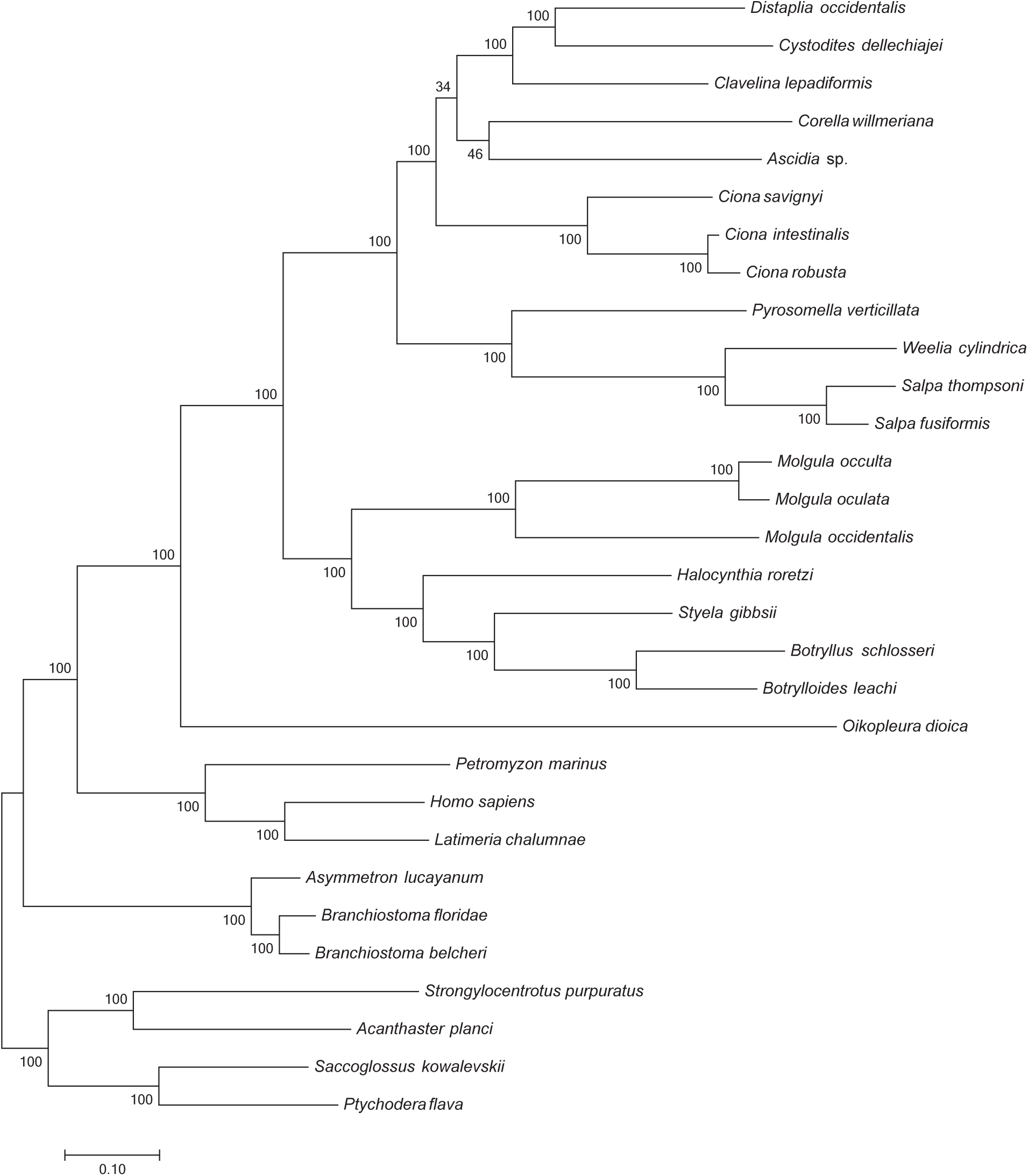
Phylogeny of Tunicata based on the RAxML analysis of the subset of the best 50 genes in the original full dataset according to RCFV retained by our pipeline after the addition of *Clavelina lepadiformis* and *Salpa thompsoni* (47 genes). Bootstrap support values are presented at each node. Scale bar represents 0.1 substitutions per site.

**Supplementary Figure 22.**
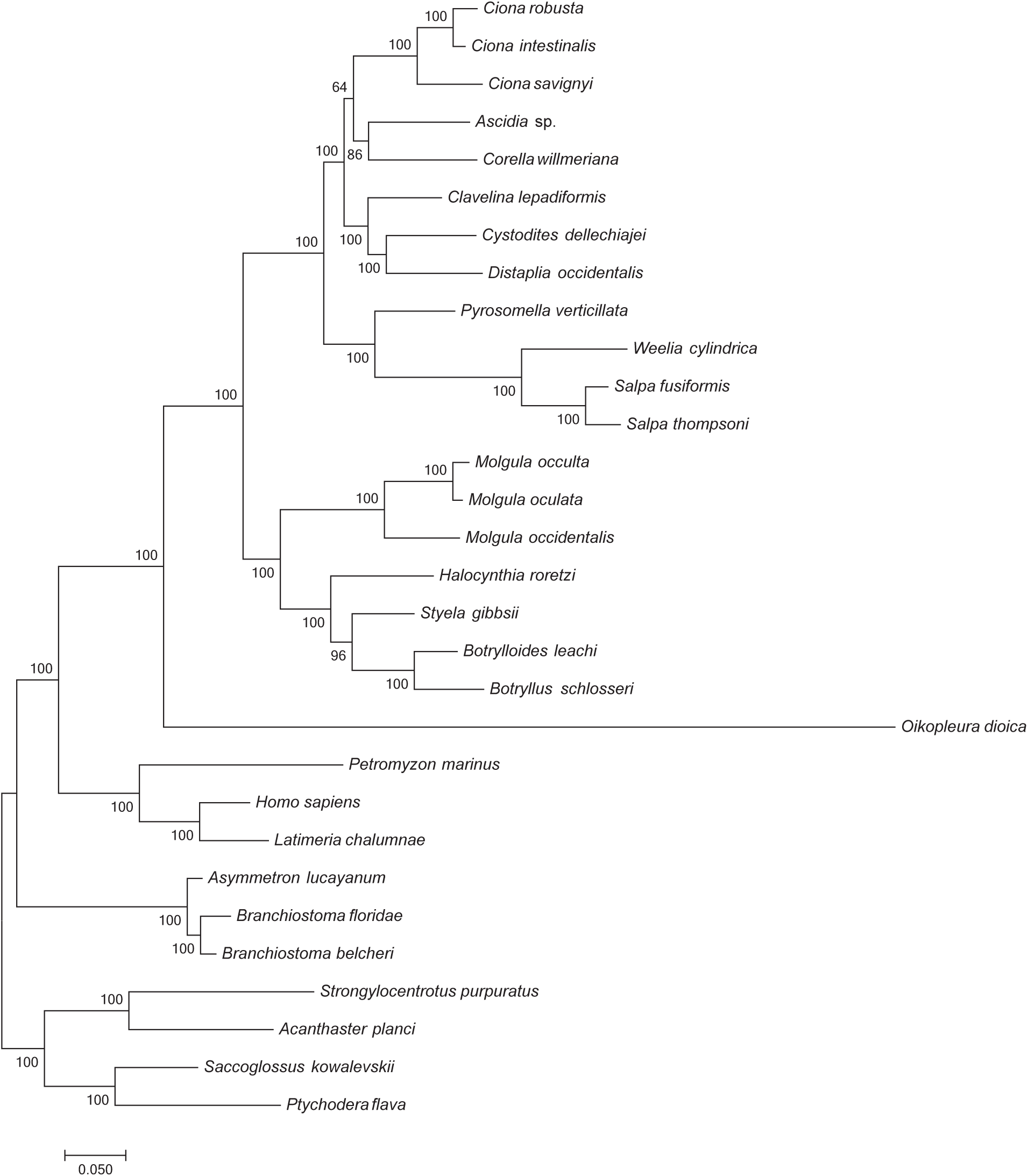
Phylogeny of Tunicata based on the RAxML analysis of the subset of the best 50 genes in the original full dataset according to LB retained by our pipeline after the addition of *Clavelina lepadiformis* and *Salpa thompsoni* (48 genes). Bootstrap support values are presented at each node. Scale bar represents 0.1 substitutions per site.

**Supplementary Figure 23.**
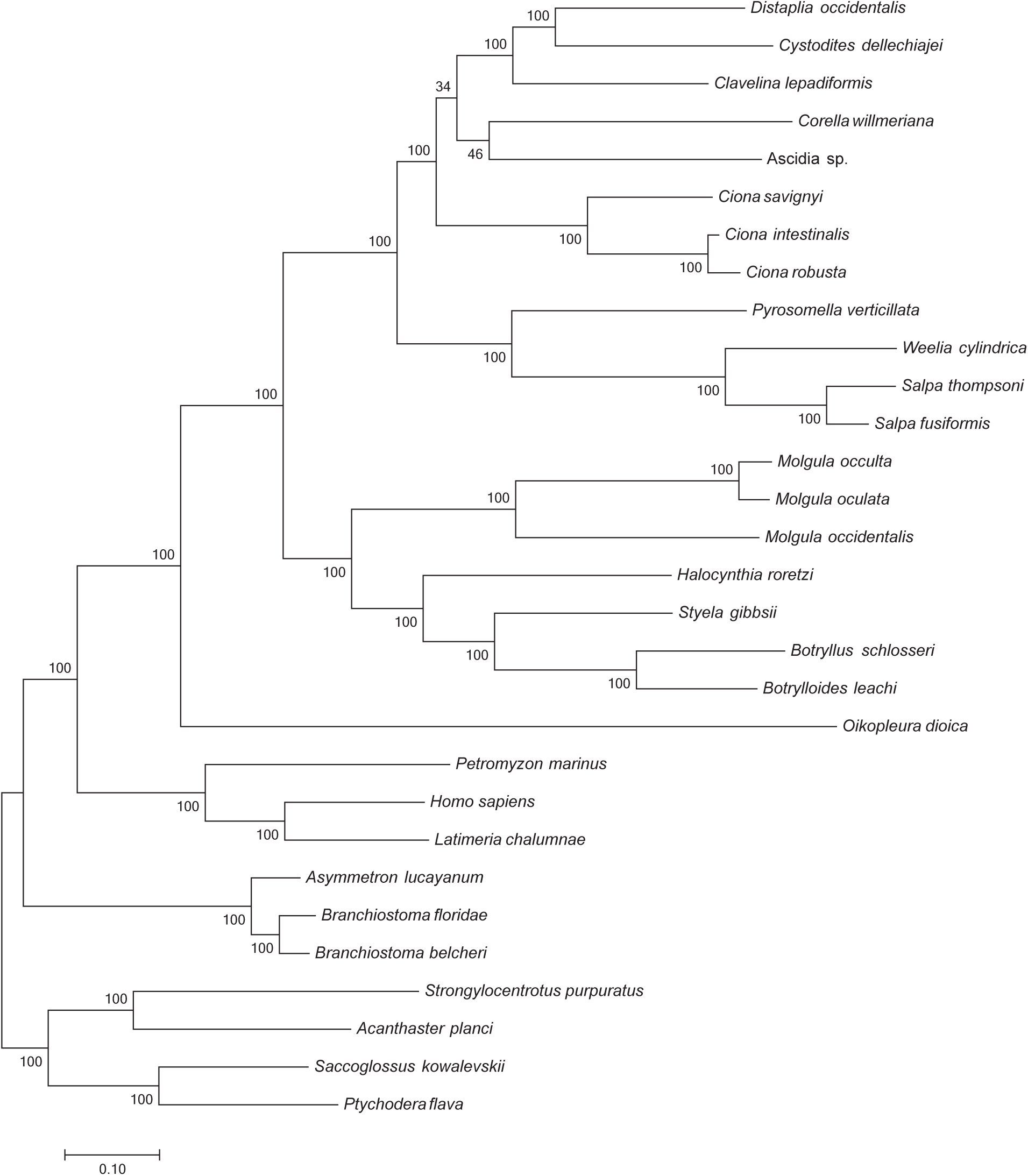
Phylogeny of Tunicata based on the RAxML analysis of all genes retained by our pipeline (788) after the addition of *Clavelina lepadiformis* and *Salpa thompsoni.* Bootstrap support values are presented at each node. Scale bar represents 0.1 substitutions per site.

**Supplementary Figure 24.**
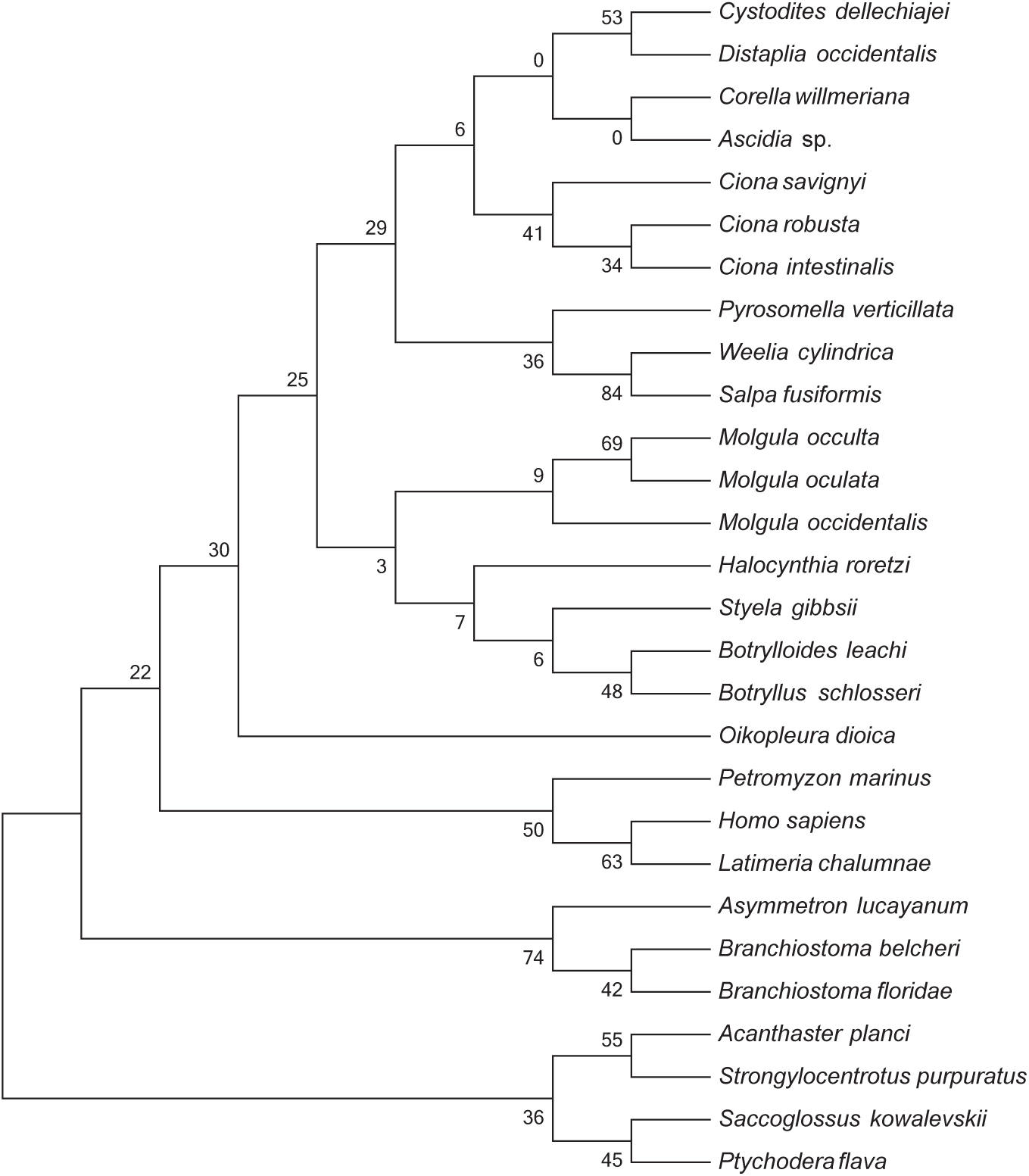
Phylogeny of Tunicata based on the RAxML analysis of all 798 genes with internode certainty scores presented at each node.

**Supplementary Figure 25.**
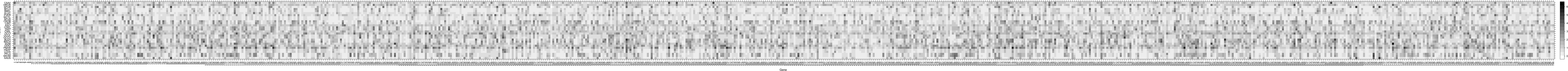
Phylo-MCOA matrix.

**Supplementary Figure 26.**
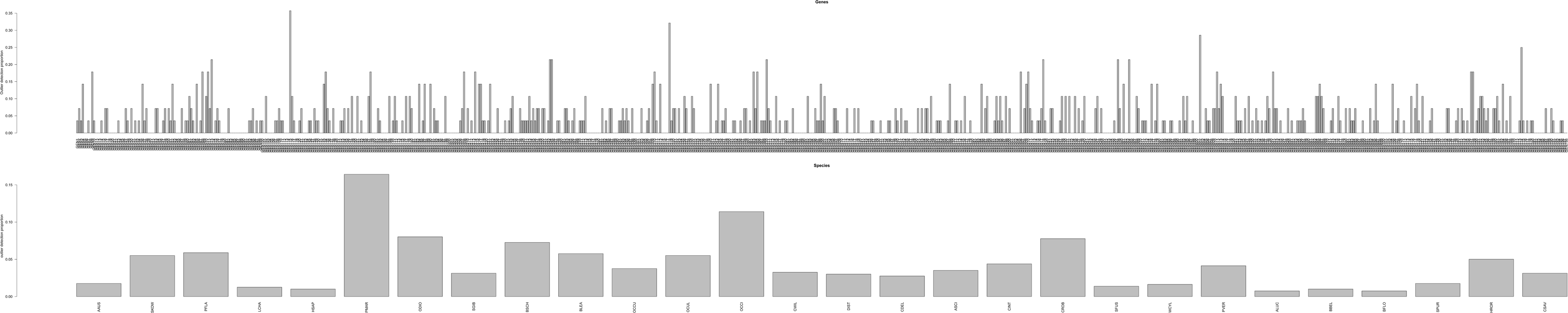
Phylo-MCOA charts.

**Supplementary Figure 27.**
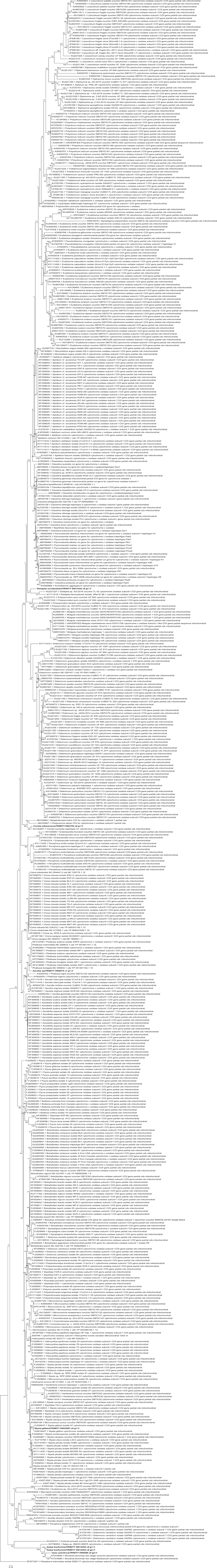
Phylogenetic analysis of COI. Bootstrap support values are presented at each node. Scale bar represents 0.1 substitutions per site. Tree presented as rooted by RAxML but should be interpreted as unrooted.

